# Picophytoplankton Implicated in Productivity and Biogeochemistry in the North Pacific Transition Zone

**DOI:** 10.1101/2025.05.29.656823

**Authors:** Rebecca S. Key, Sacha N. Coesel, Mary R. Gradoville, Rhonda L. Morales, Hanna Farnelid, Jonathan P. Zehr, E. Virginia Armbrust, Bryndan P. Durham

## Abstract

Marine phytoplankton are central to global seascapes, acting as key conduits in element cycling and oceanic food webs. Phytoplankton cell size spans several orders of magnitude (0.2 to >200 µm) and is an important trait that governs metabolism. Yet, the vast taxonomic diversity within phytoplankton size classes makes it challenging to link specific taxa to bulk community changes in productivity and elemental stoichiometry. To explore phytoplankton biogeography and biogeochemical roles in field populations, we analyzed three years of 16S and 18S rRNA gene amplicon sequencing variant (ASV) data alongside biochemical measurements across the dynamic latitudinal gradient of the North Pacific Transition Zone. We (1) identified picophytoplankton community members associated with patterns in net community production (NCP), particulate organic carbon (POC), and particulate organic nitrogen (PON), and (2) uncovered co-occurring species that may influence their growth and abundance. Multivariate linear mixed modeling revealed that occurrence of chlorophytes explained 22.6% of NCP values, followed by stramenopiles and cyanobacteria. In contrast, POC and PON spatial patterns were best explained by chlorophyte and dinoflagellate spatial patterns. Weighted co-expression network analysis further showed NCP, POC, and PON correlations with a subset of ∼40 ASVs belonging to chlorophytes, cyanobacteria, stramenopiles, haptophytes, and dinoflagellates that range in trophic strategy. Association network inference recapitulated these findings and revealed additional co-occurring phytoplankton, grazers, and heterotrophic bacteria. Together, our integrated computational analyses identified key picophytoplankton and co-occurring mixotrophs as major contributors to shaping regional biogeochemical dynamics in the North Pacific Ocean.

**Importance:** Phytoplankton mediate key biogeochemical processes in dynamic oceanic transition zones. Yet, their vast cell size range and taxonomic diversity makes it challenging to link specific taxa to bulk community changes in productivity and elemental stoichiometry. By integrating molecular and biogeochemical measurements from the North Pacific Transition Zone using combined network and multivariate modeling, we identified specific picophytoplankton strongly linked to community production and organic nutrients levels. These picophytoplankton included specific members of cyanobacteria, pelagophytes, haptophytes, and chlorophytes, and formed tight associations with several nano- and pico-sized protistan mixotrophs highlighting how top-down interactions and microbial consortia shape community structure and elemental fluxes. Our work establishes key microbial players that may control fundamental ecosystem processes like carbon and nitrogen cycling and offers a computational framework to track and identify “microbial neighborhoods” that underpin biogeochemical features of an ecosystem.

## Introduction

Photosynthetic unicellular organisms, known as phytoplankton, are the dominant players in regulating the dynamic interplay of carbon fixation, respiration, and sequestration across marine ecosystems (1). Phytoplankton cell size typically spans 0.2 µm to 1 mm, though some species can form colonies up to 1 cm in diameter. This physical size trait imposes constraints on nutrient uptake and transformation (2, 3). From a single-cell perspective, marine phytoplankton are commonly partitioned into three size classes based on cell diameter: pico- (0.2 to 2 µm), nano- (2 to 20 µm), and micro-size (20 to 200 µm) (4, 5).

In the open ocean, the majority of phytoplankton biomass is attributed to pico-sized phytoplankton, composed of cyanobacteria and the smallest eukaryotes (6). Cyanobacterial picophytoplankton primarily consist of members belonging to the genera *Prochlorococcus* (7) and *Synechococcus* (8). Meanwhile, eukaryotic members in this size range are more taxonomically diverse and consist of members spanning Haptophyta, Chlorophyta, Bacillariophyta, and Dinophyceae groups (9). Picoeukaryotic cells (∼1.5 µm diameter) are larger than *Prochlorococcus* (∼0.6 µm) and *Synechococcus* (∼1 µm), with volumes up to two orders of magnitude greater (10). This size difference, coupled with similar doubling times, helps explain why picoeukaryotes contribute up to an estimated 69% of global picophytoplankton biomass, compared to 17–39% for *Prochlorococcus* and 12–15% for *Synechococcus* (11). *Prochlorococcus* numerically dominates oligotrophic gyres, whereas *Synechococcus* and photosynthetic picoeukaryotes become more dominant in mesotrophic, coastal, and high-latitude regions. The high surface-to-volume ratios of cyanobacterial and eukaryotic picophytoplankton enhances nutrient uptake efficiency, providing a competitive advantage within their respective environments (12). Current model simulations estimate their collective contribution of roughly 58% (20% for cyanobacteria; 38% for eukaryotic picophytoplankton) to global net primary production within tropical, subtropical, and mid-latitude regions (13). In contrast, high-latitude regions with elevated nutrients and productivity are typically dominated by micro-sized phytoplankton biomass (14).

While phytoplankton distribution and productivity patterns are characterized across major ocean biomes (15), some regions exhibit greater variability in cell size, taxonomic composition, and nutrient dynamics. For example, oceanic transition zones are boundary regions between ocean biomes where surface convergence creates steep physical and chemical fronts that drive complex dynamics in phytoplankton size structure and taxonomy (16). One such zone, the North Pacific Transition Zone (NPTZ), separates the nitrogen-poor, oligotrophic North Pacific Subtropical Gyre (NPSG) from the nitrogen-replete, iron-deplete Subarctic Gyre and (17, 18). *Prochlorococcus* typically dominates the NPSG community (17), while *Synechococcus* and photosynthetic picoeukaryotes become more prevalent within the NPTZ and beyond the subarctic front (18–20). Additionally, O_2_-argon based estimates of net community production (NCP) in the NPTZ are strongly correlated with increases in nano-phytoplankton, which consist of small diatoms, dinoflagellates, and haptophytes (18). Juranek *et al.* (18) hypothesized that the observed correlation arose from a combination of increasing nutrients and grazing pressure on picophytoplankton that together facilitated co-occurrence of pico- and nano-phytoplankton. While these findings link community size structure to elevated NCP, the specific eukaryotic taxa that drive biogeochemical variability across time and space remain poorly resolved.

In this work, we investigate patterns in plankton communities and their relationships with concurrent biogeochemical features from samples collected in 2016, 2017, and 2019 during late spring and early summer along a latitudinal gradient spanning the NPSG and the NPTZ (Figure 1A). Two frontal features define gradients in the region: (1) a spatially stable salinity front (∼34.82 ppt) marking the northern edge of the NPSG, and (2) a more spatially variable chlorophyll-a front (0.15 mg m⁻³ ) that seasonally migrates from 32°N in winter to 42°N in summer, which signals an increased phytoplankton abundance and separates the southern subtropical (STZ) and northern temperate (NTZ) subregions within the NPTZ (18, 21). We used amplicon sequence variant (ASV) data derived from 18S and 16S rRNA gene sequences as proxies to infer relative abundance and community composition of microbial taxa. These data were paired with exisitng O_2_-argon-based NCP, particulate organic carbon (POC), and particulate organic nitrogen (PON) measurements (18). To investigate relationships between ASVs and biogeochemical components, we applied an integrated network and modeling strategy which combines multivariate linear mixed modeling (MLMM), weighted gene co-expression network analysis (WGCNA), and sparse inverse covariance estimation for ecological association inference (Spiec-Easi). Collectively, these approaches allowed us to assess plankton ASV contributions to NCP, POC, and PON variability along the transect and identify potential species associations that contribute to population dynamics.

**Figure 1.**
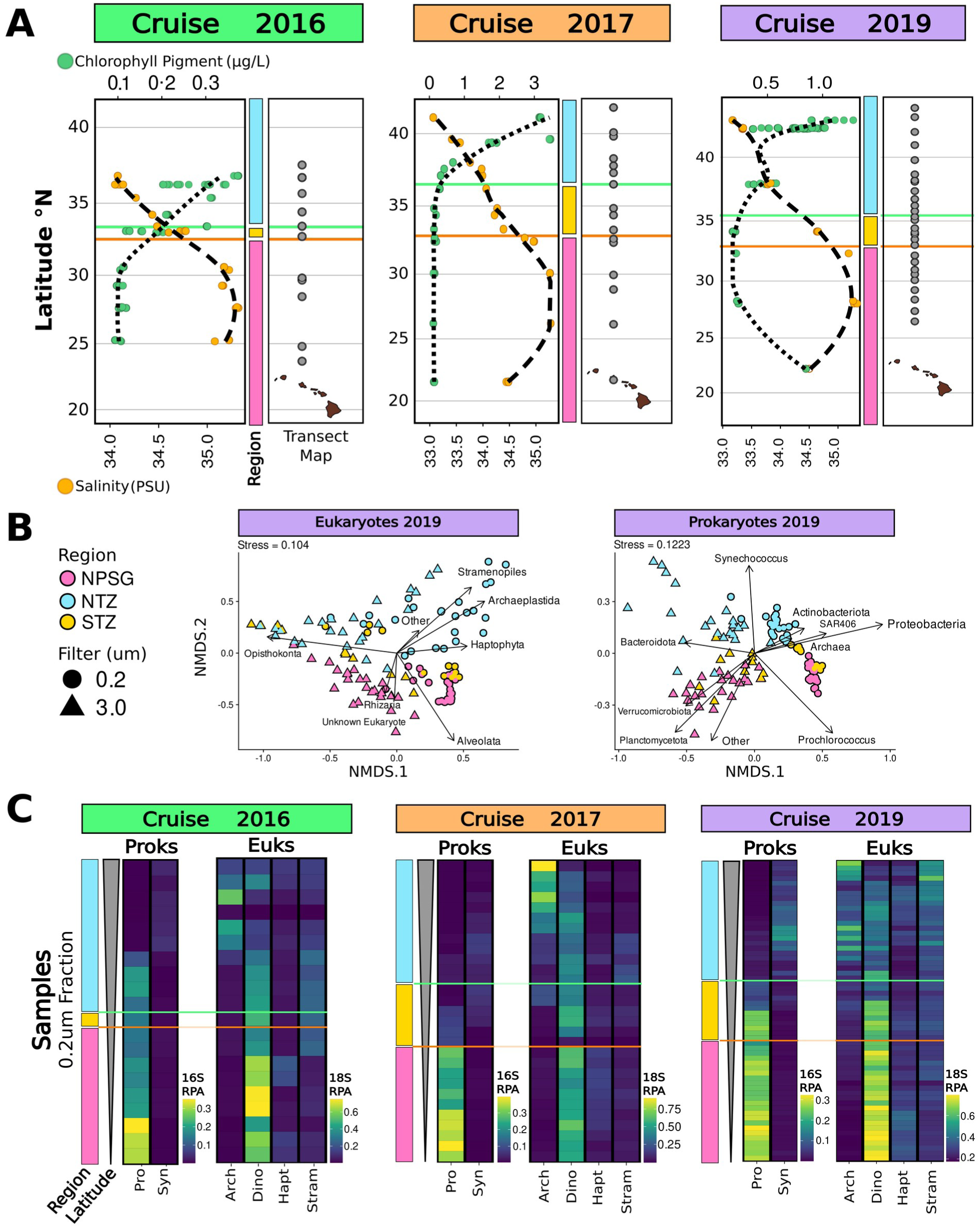
Overview of Sampling and Microbial Distributions across the Transect. **(A)** Locations of the 34.82 isohaline Salinity Front (orange line; 32.15°N–32.5°N across years) and 0.15 mg m^−3^ Chlorophyll Front (green line; 33°N–36.2°N across years) for 2016, 2017, and 2019 cruises (left), alongside ASV sampling sites (right; dark gray points). Dual X-axis plots display surface salinity (orange points) and chlorophyll (green points) at 15 m depth; dotted and dashed lines show chlorophyll and salinity trends, respectively. Color bars mark regions: North Pacific Subtropical Gyre (NSPG, pink), Southern NPTZ (STZ, yellow), and Northern NPTZ (NTZ, blue), defined by chlorophyll and salinity front positions. **(B)** NMDS of 2019 18S ASV samples collected from 0-125m depths. Circles indicate the small size fraction (0.2–3.0 µm) and triangles indicate the large size fraction (>3.0 µm), colored by region. Stress values are shown top-left. Arrows indicate correlations between samples and relative percent abundance of taxonomic groups. **(C)** Relative Percent Abundance (RPA) heatmaps of small size fraction samples collected from 0-15m depths, arranged from the bottom by increasing latitude. Columns represent phytoplankton-containing groups. Prokaryotic (Proks) and eukaryotic (Euks) amplicon datasets are displayed separately, with percent abundance of groups calculated relative to their corresponding community. The left heatmap in each survey set displays relative percent abundances for *Prochlorococcus* (Pro) and *Synechococcus* (Syn) within 16S ASVs, while the right heatmap shows Archaeplastida (Arch), Dinoflagellata (Dino), Haptophyta (Hapt), and Stramenopiles (Stram) within 18S ASVs. A color scale indicating relative percent abundance values for each dataset (16S prokaryotes and 18S eukaryotes) is provided at the bottom right. Two sample feature bars to the far left of each heatmap indicate latitude and region information salinity (orange) and chlorophyll (green) fronts as described in panel A.

## Results

### Spatial and Temporal Patterns

We examined spatiotemporal patterns of the small (0.2–3 µm) and large (>3.0 µm) size fraction 18S and 16S ASVs based on broad phylogenetic classifications representative of the most prevalent microbial groups (Figure 1). Samples spanned depths from 0 – 125 m, with the majority of samples collected from surface waters (15 m) (Figure S1). Taxonomic groups contributing <1% to total relative abundance were pooled as “Other” (Figure S2 and S3). Non-metric Multidimensional Scaling (NMDS) ordination revealed clear separation of samples based on size fraction, transect region, and water column depth across yearly surveys (Figure 1B and Table S1). For eukaryotic communities, transect region consistently explained the largest proportion of sample variation (r² = 38–47%, *P* < 0.001), followed by size fraction (r² = 17–37%, *P* < 0.001), while depth explained the least (r² = 7–13%, *P* = 0.001–0.843). Similarly, for prokaryotic communities, region explained most sample variation (r² = 46– 76%, *P* < 0.001), while filter type explained 4–12% of variation (*P* = 0.009–0.098), and depth generally had weaker or non-significant associations (r² = 0.4–41%, *P* = 0.033–0.767). NMDS also revealed the influence of eukaryotic and prokaryotic taxa on ordination placement of samples (Figure 1B and Table S2). The eukaryotic groups Opisthokonta, Alveolata, and Archaeplastida consistently accounted for a majority of variation in the ordination, with r^2^ values of 84%, 76%, and 73%, respectively (Table S2). Additional contributions came from Haptophyta (r^2^ = 23 - 39%) and Stramenopiles (r^2^ = 7 – 60%). Of the prokaryotic groups, *Prochlorococcus*, Proteobacteria, Bacteriodota, and *Synechococcus* contributed most to sample ordinates, with R^2^ values reaching 56%, 48%, 46%, and 44%, respectively (Table S2).

Plankton community composition varied markedly between size fractions and regions across the three sampling surveys (Figure 1B; Table S1 and S2). Opisthokonta (zooplankton) was a major contributor to sample separation in the NMDS ordination (Figure 1B) and was consistently dominant in the large size fraction, making up an average of 21% of the reads across samples from the three cruises (Figure S2). Alveolata were similarly represented in the small and large size fractions, reaching the highest relative abundance in the small size fraction of the NPSG (62% in 2016, 59% in 2017, and 54% in 2019). Archaeplastida, another strong contributor to eukaryotic sample ordination, was consistently enriched in the NTZ in the small size fraction, with mean relative abundance of 20% in 2016, 70% in 2017, and 29% in 2019. For prokaryotes, *Prochlorococcus* was the strongest contributor to NMDS sample ordination (Figure 1B) with consistently high relative abundance in the small size fraction in the NPSG: 30% in 2016, 37% in 2017, and 30% in 2019 (Figure S3). Proteobacteria dominated each region consistently across years, comprising over 50% of regional relative abundance within the small size fraction. *Synechococcus* was primarily concentrated in the NTZ, with relatively equal representation across both size fractions, with mean relative abundances of 2% in 2016, 5% in 2017, and 5-6% in 2019.

To more closely examine community dynamics of phytoplankton across regions, we focused on taxonomies predominantly containing photosynthetic community members (i.e., *Prochlorococcus*, *Synechococcus*, Archaeplastida, Haptophyta, Dinoflagellata within the Alveolata, and photosynthesizing Stramenopiles belonging to diatom classes Coscinodiscophyceae, Bacillariophyceae, and Mediophyceae and classes Chrysophyceae, Bolidophyceae, Dictyochophyceae, Pelagophyceae, and Pinguiophyceae). Across most groups, relative abundances of these taxonomies were consistently higher in the small size fraction (Figure S4), as compared to the large size fraction that contained relatively more Opisthokonta (Figure S2). Archaeplastida were 2- to 6-fold more enriched in the small fraction, Haptophyta 2- to 3-fold, *Prochlorococcus* 4-fold, and Stramenopiles 2- to 5-fold. In contrast, Dinoflagellata and *Synechococcus* were relatively evenly distributed across both size fractions. Examining the relative abundances in the small size fraction across the three years revealed recurring spatial patterns across major phytoplankton taxa (Figure 1C). Indeed, *Prochlorococcus* and Dinoflagellata consistently dominated the NPSG, with Haptophyta contributing to a lesser extent. In contrast, Stramenopiles, Archaeplastida, and *Synechococcus* were more prevalent in the STZ and NTZ based on ASV relative abundances.

### Persistent and Ephemeral Amplicons

We assessed ASV richness (i.e. the number of unique ASVs) within the six major phytoplankton-containing groups (Figure 2A) to determine whether the observed spatial patterns across eukaryotic and prokaryotic phytoplankton were driven by “persistent” (i.e., ASVs detected in all years) or “ephemeral” (i.e., ASVs detected in only one or two years) taxa. A majority of ASVs in both the small and large sized fraction were ephemeral, as indicated by their markedly higher contribution to richness compared to persistent ASVs. Ephemeral taxa comprised 61 - 80% of richness in 2016, 54 - 63% in 2017, and 95 - 96% in 2019. In 2017, Dinoflagellata had near-equal contribution of ephemeral and persistent ASVs (49% and 50%, respectively). *Prochlorococcus* ephemeral ASVs accounted for 81% of richness in 2016, 45% in 2017, and 95% in 2019, while *Synechoccocus* ephemeral ASVs accounted for 47%, 41%, and 93% in 2016, 2017, and 2019 respectively.

**Figure 2.**
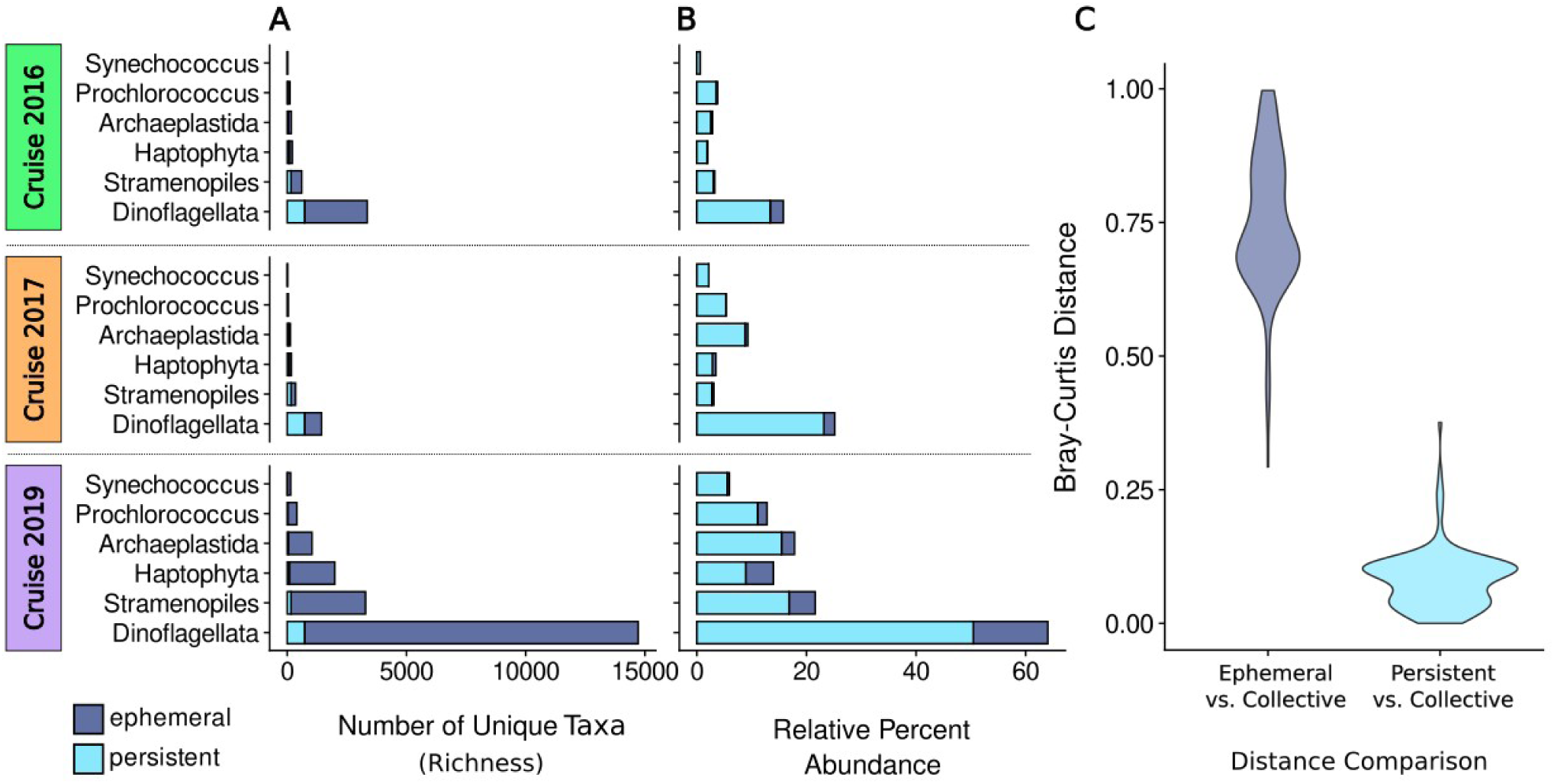
Persistent and Ephemeral ASV Comparison Across Yearly Surveys. Bar plots show the number of unique ASVs (richness) (A) and total relative percent abundance of ASVs (**B**) across six phytoplankton-containing groups for 2016, 2017, and 2019 cruises. Richness is divided into persistent (cyan) and ephemeral (dark blue) categories, with persistent ASVs defined as those present in all three cruises and ephemeral defined as those identified in < 3 cruises. Samples collected from all depths are considered. **(C)** Violin plots show Bray-Curtis distances between ephemeral (left) and persistent (right) group compositions and the overall community across all years. Each distribution represents the dissimilarity between ephemeral or persistent groups and the overall community, with higher distance values indicating greater dissimilarity from the overall relative percent abundance.

In contrast to their generally low contribution to total richness, persistent ASVs contributed substantially to total relative abundance (Figure 2B). Persistent ASVs composed over 77% of the overall relative abundance each year across both eukaryotic and prokaryotic groups. Cyanobacteria had the highest contributions, with persistent *Synechococcus* and *Prochlorococcus* ASVs accounting for 93 - 99% and 86 - 99% of overall relative abundance, respectively. A mean Bray-Curtis dissimilarity distance of 0.75 (median 0.72) distinguished the relative abundance of ephemeral ASVs and the overall dataset of ephemeral and persistent ASVs (Figure 2C). On the other hand, persistent ASVs exhibited a mean value of 0.08 (median 0.09) when compared to the overall dataset. Thus, persistent ASV distribution patterns more closely reflect the distribution patterns of the overall community, and we proceeded to use persistent taxa in subsequent network and modeling strategies.

### Phytoplankton Distribution and Covariation with Biogeochemical Trends

We evaluated which phytoplankton-containing groups were most closely aligned with previously published (18) concurrent measurements of net community production (NCP), particulate organic carbon (POC), and particulate organic nitrogen (PON) (Figure 3A). Mean NCP, POC, and PON levels were highest within the NTZ (39.6 mmol O_2_ m^-2^ d^-1^, 18.2 *μ*mol C L^-1^, and 2.7 *μ*mol N L^-1^, respectively) followed by the STZ (10.7 O_2_ m^-2^ d^-1^ , 6.7 *μ*mol C L^-1^, and 1.3, respectively), with lowest values in the NPSG (8.4 m^-2^ d^-1^ , 2.7 *μ*mol C L^-1^, and 0.4 *μ*mol N L^-1^, respectively) (Figure 3A).

**Figure 3.**
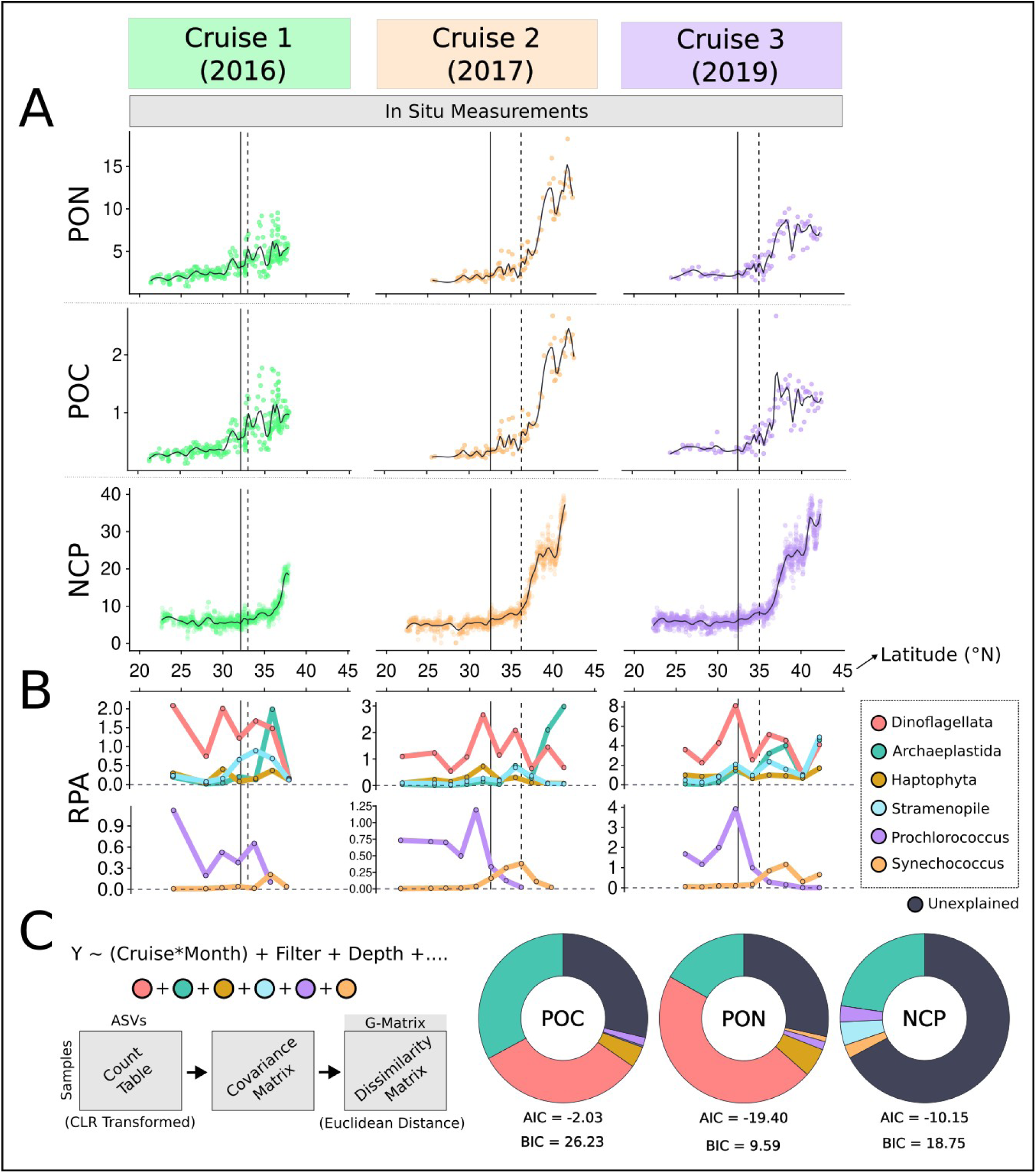
Microbial Plankton Contributions to Spatio-Temporal Variations in Biochemical Parameters. **(A)** Dot plots show latitudinal patterns of surface POC, PON, and NCP (0-15m depth) from 2016 (green), 2017 (orange) and 2019 (purple) surveys. Solid and dashed lines denote the salinity front (isohaline 34.82) and the chlorophyll front (0.15 mg m^−3^), respectively. Black solid curve lines represent the fitted trend of the data. **(B)** Relative Percent Abundance (RPA) of eukaryotic (top) and cyanobacterial (bottom) small-size fraction phytoplankton across 2° latitude bins, highlighting changes in group dominance alongside biogeochemical patterns in panel A. Samples collected from all depths are considered. **(C)** Multivariate Linear Mixed Model formula (left) illustrating fixed effects (year interacting with month, filter size, and water column depth) and random variables (colored circles; phytoplankton-containing groups), with both size fractions included during modeling. Filter size categories include: 0.2µm and 3µm. Depth categories include 0-15m, 45-75m, and 90-125m. Each response variable (Y) represents a biochemical feature (POC, PON, or NCP). Below, a schematic shows how G-matrices were built from ASVs grouped by each phytoplankton group. Donut plots (right) display the variance per biochemical feature explained by each phytoplankton group: Dinoflagellata (red), Archaeplastida (green), Haptophyta (yellow), Stramenopiles (blue), *Prochlorococcus* (orange), and *Synechococcus* (purple). Black segments indicate unexplained variance. Akaike Information Criterion (AIC) and Bayesian Information Criterion (BIC) scores below each plot assess model fit.

We performed multivariate linear mixed modeling (MLMM) to assess the degree of covariation between persistent cyanobacterial and eukaryotic phytoplankton community structure and patterns in NCP, POC, and PON across the latitudinal transect (Figure 3C). Dissimilarity-based G matrices, built from ASV-level data within each major phytoplankton-containing group, preserved fine-scale taxonomic resolution while capturing covariation with NCP, POC, and PON. Archaeplastida and Dinoflagellata were the top contributors to PON variability across the transect, explaining 17% and 47% of the variance, respectively (AIC: -19.4; BIC: +9.6). Minimal covariation was found for Haptophyta (5.5%) and *Prochlorococcus* (1.6%), with 28% of the variance unexplained potentially due to unmeasured environmental factors or stochastic processes. Archaeplastida and Dinoflagellata each explained ∼32% of the POC variance (AIC: –2.03; BIC: +26.2). Haptophyta (4%) and *Synechococcus* (2%) contributed minimally, while 29% of POC patterns remained unexplained. Archaeplastida showed the highest covariation with NCP (23%; AIC: –10.1; BIC: +18.8), followed by Stramenopiles (4.6%), *Prochlorococcus* (3%), and *Synechococcus* (3%), with 67% of NCP variance unexplained. While Archaeplastida distributions positively covaried with PON and POC values, Dinoflagellata had inverse relationships, suggesting the two groups play contrasting roles in POC and PON dynamics (Figure 3B).

### Community-Wide Correlation of Microbial Plankton and Biochemical Variability

Weighted gene correlation network analysis (WGCNA) was used to determine whether specific persistent phytoplankton ASVs were correlated with the different environmental variables. The resulting network was organized into 13 distinct modules (clusters), with 11 forming a densely connected central core and two (yellow and purple) forming smaller, more isolated clusters (Figure 4A). NCP, POC, and PON were significantly associated with ASVs in the yellow (r = 0.7, 0.7, and 0.6, respectively; P < 0.001) and purple clusters (r = 0.6, 0.3, and 0.3; P < 0.001), with all three variables assigned to the yellow cluster (Figure 4A). Regenerating the WGCNA network at varying powers (powers ranging from 3 to 8) confirmed that ASVs found within yellow and purple clusters remained consistent (Figure S5). In contrast to yellow and purple clusters, several central core clusters exhibited predominately negative correlations with biogeochemical variables (r = -0.02 to -0.5, P < 0.001), with the red cluster being the most negatively correlated. Other core clusters like the lime cluster showed weak and non-significant correlations with POC and PON (r = 0.04, 0.05; P = 0.4–0.5, respectively), while the clay and black clusters displayed weak positive associations with NCP (r = 0.2, 0.1; P = 0.004, 0.051, respectively). These results suggest that the ASVs in the yellow and purple clusters may play an important role in shaping NCP, POC, and PON dynamics.

**Figure 4.**
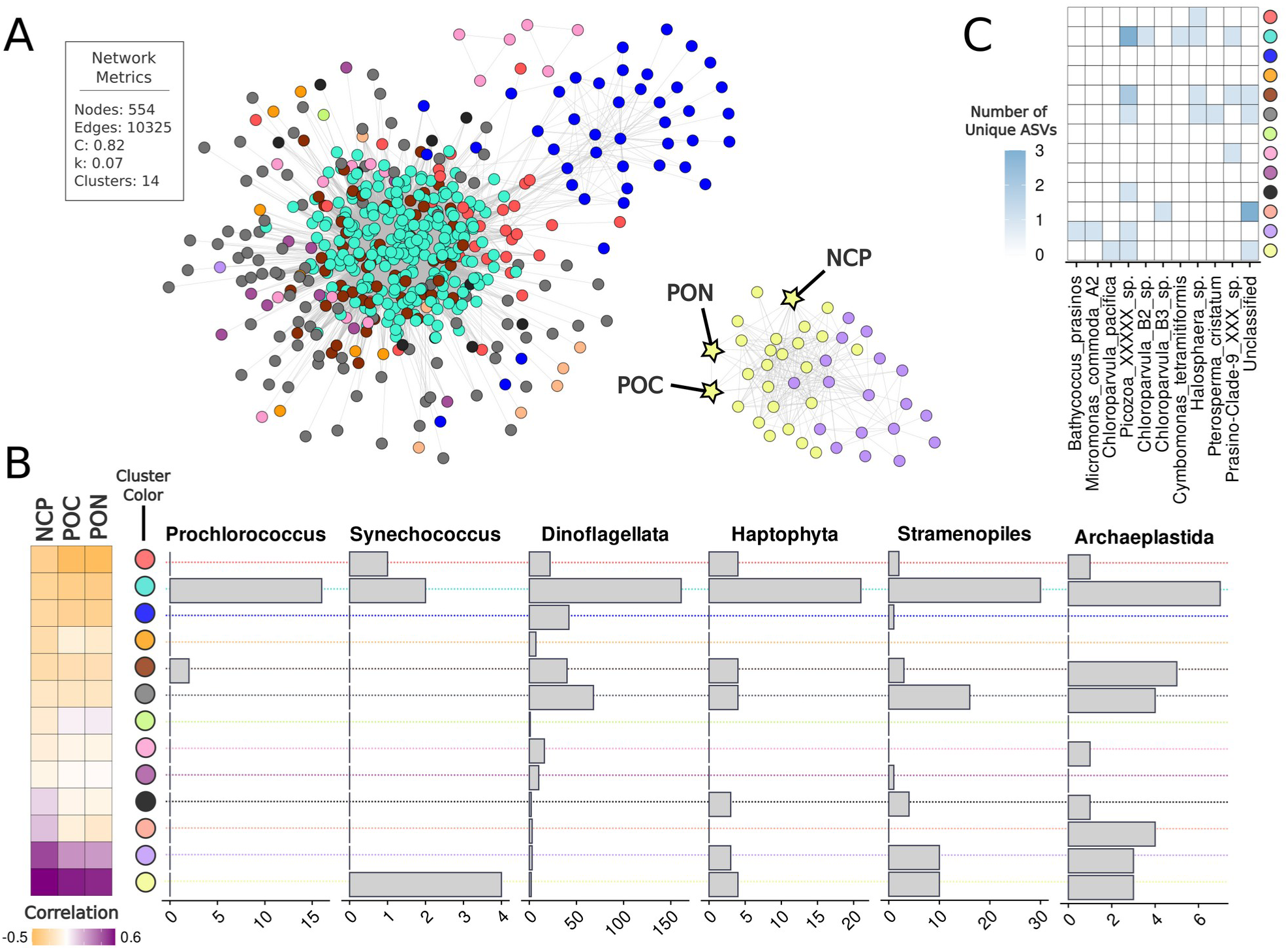
WGCNA of Phytoplankton-Containing Communities and their Association with Biochemical Measurements. **(A)** Network visualization showing ASV modules identified through weighted gene correlation network analysis (WGCNA). Nodes are colored by module (cluster of ASVs) that exhibit coinciding spatial patterns across the transect. Nodes represent individual persistent ASVs, while edges represent the potential interactions existing between them. The biochemical measurement nodes NCP, POC, and PON are represented by yellow stars positioned among their most strongly correlated modules. The network consisted of 554 nodes and 10,325 edges, with a mean clustering coefficient (C) of 0.8 and network connectivity (k; density) of 0.07, organized into 13 clusters (see inset). Samples with available NCP, POC, and PON measurements were used in network construction. **(B)** Correlation plot (far left) and bar plots (right) showing module relationship to biochemical measurements and taxonomic composition of each cluster, respectively. Cluster colors are provided between the correlation and taxonomy plots and correspond to the WGCNA network in Panel A. Taxonomic bar plots show the number of unique ASVs within each major phytoplankton-containing group (*Prochlorococcus, Synechococcus,* Dinoflagellata, Haptophyta, Stramenopiles, Archaeplastida). **(C)** A richness heatmap showing the taxonomic distribution of key Archaeplastida species across each network cluster in Panel A. Colors represent the number of unique ASVs for each species, from zero (white) to three (blue). Network cluster colors are provided to the right of the heatmap.

The yellow cluster contained 10 ASVs from Stramenopiles, three from Archaeplastida, four from Haptophyta, two from Dinoflagellata, and four from *Synechococcus* (Figure 4B and Table S3). Archaeplastida members included *Chloroparvula pacifica*, an unclassified chlorophyte, and an unclassified Picozoa (Figure 4C). Although classified within Archaeplastida, Picozoa are currently considered heterotrophic and distinct from chlorophytes (22). The cluster also featured other heterotrophic and mixotrophic protists, such as *Chrysochromulina* sp. (Haptophyta), MAST-1A, -1B, - 1C and -7A sp. (Stramenopiles), and a Parmales bolidophyte (Stramenopiles). Additional members were the parasitic dinoflagellate Syndiniales, haptophyte *Phaeocystis antarctica*, and stramenopile *Pseudochattonella*.

In the purple cluster, there were 10 ASVs from Stramenopiles, three from Archaeplastida, three from Haptophyta, and three from Dinoflagellata (Table S3). Archaeplastida members included *Bathycoccus prasinos, Micromonas commoda A2*, and another unclassified Picozoa (Figure 4C). Like the yellow cluster, several ASVs belonged to *Chrysochromulina* sp., Syndiniales, and Parmales (*Triparma pacifica*). Other notable members included the haptophyte *Phaeocystis pouchetii* and stramenopiles such as *Pelagomonas calceolata, Aureococcus anophagefferens* (pelagophytes), *Dictyocha speculum* (silicoflagellate), *Brockmanniella brockmannii* (diatom), and a member of the MAST-2D lineage.

### Key Species Co-occurrence Patterns

We used Spiec-Easi network analysis to identify taxa that displayed co-occurrence patterns and thus may potentially influence NCP, POC, and PON dynamics through metabolic interactions. The Spiec-Easi network included persistent ASVs from the six phytoplankton-containing groups, as well as all persistent 16S ASVs. This resulted in a moderately connected structure with 1,885 nodes and 37,350 edges averaging 40 neighbors per ASV (Figure 5). Of these nodes, 1,006 were eukaryote ASVs, and 879 were prokaryote ASVs. As an internal validation, we examined whether a known association was recovered from this network: specifically, the symbiosis between *Braarudosphaera* and the nitrogen-fixing UCYN-A, shown to be an early-stage organelle, or nitroplast (23–25). Their co-occurrence within the network aligned with these prior studies (Figure 5). Notably, the network was fully connected with no disconnected subnetworks. The *Chloroparvula pacifica* ASV from the WGCNA yellow cluster and the unclassified Picozoa ASV from the purple cluster were among the ASVs with highest connectivity (i.e. out-degree) for eukaryotes (Figure 5C; Table S4), while ASVs from Alphaproteobacteria, Bacteroidia, and Actinobacteria contained the highest out-degree connections within the heterotrophic bacterial community (Table S4).

**Figure 5.**
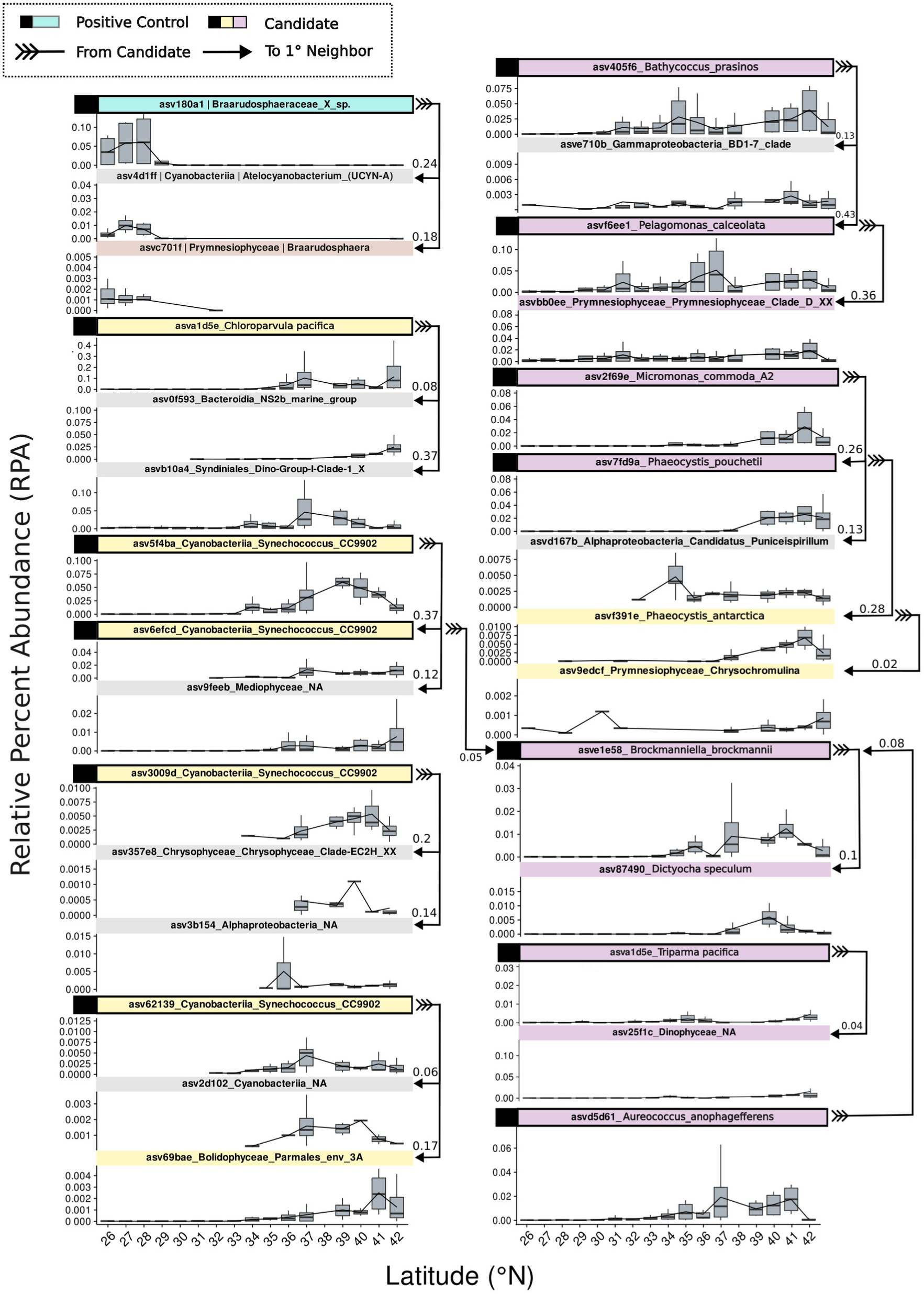
In-Situ Distributions of Yellow and Purple Cluster ASVs with their Top 1^°^ Spiec-Easi Neighbors. Relative percent abundance distributions in 2019 of yellow and purple ASVs (i.e. Candidates) with full binomial species-level classifications (i.e., both genus and specific epithet), along with their top positively weighted first-degree neighbors from persistent phytoplankton and prokaryotic ASVs. Facet labels display each ASV’s unique ID, taxonomic classification, and WGCNA cluster assignment by color. Neighbors with no cluster assignment are shown in gray. Arrows illustrate first-degree connections, originating from the candidate (feathered end) and pointing to its neighbor. Latitude is plotted on the x-axis, and relative percent abundance is shown on the y-axis. As internal validation, a known symbiosis between *Braarudosphaera* (the most abundant ASV from the family *Braarudosphaeraceae* assigned to the teal cluster) and the nitrogen-fixing cyanobacterium UCYN-A (*Candidatus* Atelocyanobacterium thalassa) was detected (top-left of the panel). Samples from all depths were used in Speic-Easi network construction.

We analyzed weights and cluster assignments for first-degree (1°) neighbors (i.e. ASVs connected directly by one network edge) for ASVs within both yellow and purple WGCNA clusters that positively correlated to NCP, POC, and PON measurements (Fig. S6). Association weights ranged from –0.04 to 0.44 (mean = 0.07), with largest absolute weight values indicating strongest associations. Positive associations spanned 0.0005 to 0.440, while negative associations ranged from –0.04 to –0.001 (Fig. S6A). Positively weighted 1° neighbors tended to belong to the same yellow and purple WGCNA clusters (Fig. S6B). In contrast, negatively weighted neighbors were primarily housed in WGCNA clusters that negatively correlated with NCP, POC, and PON measurements, such as the red and cyan clusters (Figures 4B and S6B). Transect relative abundance profiles for positive neighbor connections to yellow and purple candidates with complete binomial species classifications (i.e. genus and species epithet) are highlighted in Figure 5. We specifically focused on neighbors with the highest connectivity (i.e. weight). All positive yellow and purple WGCNA neighbor-candidate connections are in Table S5.

From the yellow cluster, the *Chloroparvula pacifica* ASV showed highest positive connection to a Syndiniales Dino-Group-1-Clade-1 ASV (weight = 0.37) among eukaryotic associates and an NS2B Bacteroidia NS2b Marine Group ASV (weight = 0.09) among prokaryotic associates (Figure 5C). Four *Synechococcus sp.* CC9902 ASVs were also found within the yellow cluster. One with the identifier ‘5f4ba’ exhibited highest connection with an unknown diatom species from Mediophyceae (0.12) and with another yellow-cluster *Synechococcus* sp. CC9902 ASV (0.37). This second *Synechococcus* ASV showed its highest-weighted connection to a purple-cluster diatom *Brockmanniella brockmannii* (0.05). The remaining two *Synechococcus* sp. CC9902 ASVs showed highest-weighted eukaryotic connections to an unclassified Chrysophyceae ASV (0.2) and the yellow-cluster Parmales ASV (0.17) and prokaryotic associations to an unclassified Alphaproteobacteria (0.14) and Cyanobacteria (0.06), respectively (Figure 5C). The same Parmales ASV was also connected to a MAST-1A ASV, also from the yellow cluster (Table S5). We also found the *P. antarctica* ASV connected to a yellow-cluster *Chrysochromulina* ASV (0.3) (Figure 5C) which shared an edge with the same MAST-1A ASV (0.18) mentioned above (Table S5). This MAST-1A ASV itself neighbored two additional yellow-cluster stramenopiles: a Parmales_env_3A ASV (0.11) and a MAST-7A ASV (0.05). No positive prokaryotic neighbors were detected for the mentioned Parmales, *P. antarctica*, *Chrysochromulina,* MAST and Bolidophyceae ASVs.

For purple-cluster Archaeplastida members, the *Bathycoccus prasinos* ASV showed the highest positively weighted connection with another purple-cluster member *Pelagomonas calceolata* (0.44) and a gammaproteobacterium ASV from the BD1-7 clade (0.13) (Figure 5C). In contrast, *P. calceolata* itself had no prokaryotic associates, instead linking to an unclassified Prymnesiophyceae Clade D ASV also assigned to the purple cluster (0.36). For *Micromonas commoda*, top associations included the purple-cluster haptophyte *Pelagomonas pouchetii* (0.27) and a SAR116 clade ASV (0.13). *Triparma pacifica* showed highest eukaryotic connection to an unclassified Dinophyceae ASV (0.04) and lacked prokaryotic associations. The purple-cluster diatom *B. brockmannii* was associated with *Dityocha speculum* (0.10) and *Aureococcus anophagefferens* (0.08), both of which belonged to the purple cluster, but showed no positive links to prokaryotic ASVs. Lastly, *P. pouchetii* exhibited its highest-weighted connection with relative *P. antarctica*, which belonged to the yellow cluster (0.28).

## Discussion

Our spatiotemporal study revealed distinct shifts in phytoplankton cell size composition and community structure across the natural physio-chemical gradient of the NPTZ (Fig. 1C; Fig. S2 and S3). In the southern region of the transect, *Prochlorococcus* (pico-plankton) and Dinoflagellata (nano- and micro-plankton) relative abundances were the highest but then declined just north of the NPSG, a trend consistent with previous studies that analyzed biomass estimates and metatranscriptomic data of major taxonomic groups (17, 23). Within the more productive NTZ, *Synechococcus* (pico-), Archaeplastida (pico-), and Stramenopiles (pico- to micro-plankton) increased in relative percent abundances. Our study revealed that persistent ASVs (those detected in all three years) dominated the total relative abundance and reflected overall community patterns across all cruises (Figure 2). This suggests a stable “core” set of taxa that control community structure and biogeochemistry in the NPTZ. By linking compositional shifts of these core taxa with concurrent biochemical trends, we identified specific plankton taxa, particularly pico-sized groups, that likely shape elemental stoichiometry and ecosystem productivity. Furthermore, comparing another network approach more tailored for compositional data (Spiec-Easi), rather than phenotype–trait relationships (WGCNA), revealed shared overlap and insight into putative trophic interactions to explore further.

### Microbial Plankton Composition and Stoichiometry

Stoichiometry measurements from the North Pacific have demonstrated that the NPSG exhibit elevated C:N ratios relative to Redfield, whereas the NPTZ typically display lower ratios (26). This study builds on the Redfield Ratio theory (27) by examining how phytoplankton community composition relates to POC and PON levels across a latitudinal gradient. While the Redfield Ratio (106:16:1) has historically been used to describe marine particulate matter composition, numerous studies have demonstrated that elemental stoichiometry is highly variable across oceanic regions, particularly in response to environmental gradients (26–28). Whether these regional patterns are primarily driven by taxonomic composition or environmental constraints is debated. Some studies argue that oceanographic conditions dictate stoichiometric variability (28, 29), while others suggest that taxonomic differences in macromolecular allocation play a critical role (30, 31).

Previous work has shown that dinoflagellates exhibit higher C:N ratios, likely reflecting an increased allocation to carbon-enriched macromolecules, such as energy-storage lipids (31). This trait likely enhances their ability to thrive in oligotrophic environments where nitrogen availability is limited, a pattern consistent with their high relative abundance in the NPSG (Figure 3A). Conversely, Archaeplastida display lower C:N ratios compared to Dinophyceae members (31), suggesting a more balanced allocation of carbon and nitrogen. This stoichiometric balance likely contributes to the success and increase in relative abundance of Archaeplastida taxa in nitrogen-enriched regions of the NPTZ (Figure 1C). Our findings align with previous arguments by Quigg et al. and Garcia et al., who proposed that differences in elemental stoichiometry among phytoplankton groups arise from evolutionary trade-offs in macromolecular allocation, particularly under varying nutrient regimes, which in turn shape their biogeographic distributions (30, 31). We observed distinct shifts in community structure corresponding to changes in POC and PON, particularly for Archaeplastida and Dinoflagellata, that support this framework (Figure 3).

### Chlorophyta Populations and NCP

Archaeplastida were consistently enriched in the NTZ (Figure 1C) and dominated by a few persistent ASVs (Figure 2A-B). Members of Archaeplastida exhibited strong covariation with POC, PON, and NCP based on MLMM (Figure 3C), and the WGCNA identified specific chlorophyte ASVs closely aligned with NCP, such as *B. prasinos, C. pacifica, and M. commoda A2* (Figure 4C). The picoeukaryotes, *Bathycoccus* and *Micromonas* dominate nutrient-rich coastal regions and sporadically bloom under favorable conditions (6, 32–34). *Bathycoccus* has also been observed in deeper epipelagic zones where light and nutrient availability are more limited (35), and *Micromonas* is an important member of eukaryotic picophytoplankton communities in the Arctic, Atlantic temperate waters, and various coastal environments (36). Our findings further highlight the importance of chlorophyte picoeukaryotes to open-ocean carbon and nutrient cycling.

Recent advances in molecular-based taxonomic resolution have expanded our understanding of open-ocean *Chlorophyta* members, particularly through the identification of chlorophyte clade VII, which dominates surface photic zones in oligotrophic marine environments (33, 37). This clade was recently revised with the formal description of two novel classes: Chloropicophyceae and Picocystophyceae, which contain the newly described genera *Chloropicon* and *Chloroparvula*. Notably, we identified *C. pacifica* as a persistent ASV in the NTZ that exhibited strong covariation with NCP measurements, suggesting its role in regional carbon dynamics (Figure 4C). Our findings highlight chlorophytes as key players in regional carbon and nitrogen dynamics and a need for continued investigation of recently resolved *Chlorophyta* picophytoplankton groups.

### Protistan Mixotrophs and Grazers

We identified several mixotrophic and heterotrophic protist ASVs coinciding with picophytoplankton communities based on WGCNA clustering and Spiec-Easi-derived associations. Unclassified Picozoa were detected in the WGCNA yellow and purple clusters, which also included *Synechococcus* and autotrophic chlorophytes. Picozoa are currently uncultured members of the Archaeoplastida, deemed to lack plastids, and presumed to have a heterotrophic lifestyle (22, 38). Genomic evidence has revealed viral-origin DNA within their cells, raising the possibility of viral particle ingestion (22, 39). The co-occurrence of Picozoa members with *Synechococcus* and other pico-sized phytoplankton in the transition zone may indicate an indirect, positive association. For instance, Picozoa may prey on viruses that target *Synechococcus* or picoeukaryotic phytoplankton, potentially modulating host populations via viral grazing (40). Alternatively, they may consume microbial antagonists that negatively affect picophytoplankton. Early reports of eukaryotic algal viruses described large DNA viruses infecting marine chlorophytes, particularly *Bathycoccus* and *Micromonas* (41–43). If chlorophyte populations are regulated by viruses or microbial antagonists, such as the Spiec-Easi neighbors Alphaproteobacteria (*Micromonas*-associated) and Gammaproteobacteria (*Bathycoccus*-associated) (Figure 5), it is possible that Picozoa could indirectly relieve these pressures on picophytoplankton populations.

We also observed several associations that may reflect direct trophic interactions, such as grazing by larger protists. Four *Synechococcus* ASVs within the WGCNA yellow cluster co-occurred with members of heterotrophic MAST lineages (MAST-1, MAST-7), mixotrophic *Chrysochromulina* sp. (Haptophyta), and the Parmales group. The SpiecEasi network additionally detected association between Parmales and *Synechococcus* ASVs (Figure 5). Prior studies have demonstrated that *Synechococcus* populations are strongly influenced by top-down controls from plankton grazers, including both heterotrophic and mixotrophic protists. For example, recent genomic analyses suggest that Parmales may exhibit phago-mixotrophy, combining photosynthesis with the ingestion of prey to better thrive under nutrient-limited conditions (44). Additionally, a past study in the East China Sea during the summer season demonstrated that MAST grazers strongly correlated with *Synechococcus* populations, with experimental validation confirming their ingestion of *Synechococcus* in culture (45).

While MAST members were not the strongest eukaryotic neighbors to any *Synechococcus* ASVs, they exhibited the strongest weighted association with purple cluster members *Chrysochromulina* and Parmales, which in turn connected to *Synechococcus*, suggesting an indirect influence on its population dynamics. These patterns highlight the complexity of picoplankton interactions within the NPTZ and underscore the need for future studies to disentangle the nature and directionality of these putative associations.

## Conclusion

Microbial ‘neighborhoods’ shape ecosystem dynamics through ecological associations, wherein metabolic coupling, niche construction, and trophic interactions form an emergent, self-sustaining unit that impacts broader ecosystem dynamics (46–48). Our integrated modeling and co-occurrence network strategies proved useful to identify key neighborhood members in the NPTZ. Unlike direct interaction studies, network-based and systematic modeling approaches capture emergent patterns of microbial co-existence found in nature, revealing hidden ecological roles that shape elemental biogeochemical cycles. We recapitulated the well-documented association between *Braarudosphaera* and its nitrogen-fixing symbiont/organelle UCYN-A (23, 49), a relationship currently considered obligate. We also identified a chlorophyte-centric neighborhood that likely underlies enhanced NCP and altered stoichiometry in the NPTZ, and interestingly, may be directly or indirectly controlled by grazing. These observed associations highlight the use of network analysis to detect a range of complex associations, some of which may operate through direct and indirect ecological mechanisms. As the field of microbial ecology advances, statistical modeling and network-based co-occurrence analyses will be essential for unraveling the complexity of microbial interactions, offering a systems-level approach to identifying both emergent symbiotic relationships and broader ecosystem dynamics.

## Materials and Methods

### Amplicon Collection and Sequencing

Seawater samples for 16S and 18S rRNA gene sequencing were collected during three cruises: Gradients 1 (April 19–May 5, 2016; 23°N to 37°N), Gradients 2 (May 25–June 13, 2017; 21°N to 41°N), and Gradients 3 (April 9–30, 2019; 26°N to 42°N), all along the ∼158°W longitude. A total of 48 samples were collected in 2016, 72 in 2017, and 192 in 2019. Samples were collected via CTD rosettes or shipboard surface-intake systems. Sampling depth ranged from 0-110 m in the 2016 survey, 12-15 m for 2017, and 0-125 m for 2019. For each sampling site with a unique latitude, depth, and time, 1-3 replicates were collected per size fraction (Dataset S1). Salinity and chlorophyll fronts delineating NPTZ regions were defined following Juranek *et al.* (18).

A peristaltic pump was used to sequentially filter cells onto 3.0-μm pore size 25 mm polyethersulfone membranes (Sterlitech, Kent), and 0.2-μm pore size 25 mm Supor membranes (Pall Corporation, New York). Filters were immediately flash frozen in liquid nitrogen and subsequently stored at -80°C until processing. Genomic DNA was extracted following protocols from Gradoville *et al.* (21). In brief, samples were processed using Dneasy Plant Mini Kits following manufacturer protocols (Qiagen, Venlo, the Netherlands). Additional steps included three freeze-thaw cycles, two minutes of bead-beating, and a Proteinase K treatment, as described by Moisander *et al.* (50). The V4 region of the 16S rRNA gene was amplified using primers 515F/806R (51), and the V4–V5 region of 18S rRNA gene using 566F/1200R (52). These amplified genetic loci regions were then barcoded, quantified, pooled, and sequenced using 250 bp paired-end reads on an Illumina MiSeq.

### Amplicon Processing & Community Analysis

ASVs were processed using QIIME2 (53). Sequences were quality-controlled, trimmed, merged, and annotated as described in the Supplemental Methods. ASV counts were rarefied (Table S6) prior to calculating relative percent abundance values. 16S and 18S ASVs were annotated against the SILVA database (54) and PR2 database v5.1 (55, 56), respectively and further categorized as either *persistent* (present in all three cruises) or *ephemeral* (present in only one or two cruises). NMDS was performed using *vegan* v2.6-4 (57). Details can be found in the Supplemental Methods.

### Physio- and Biochemical Data Integrations

Physiochemical measurements (seawater surface temperature and salinity) and biochemical measurements (POC, PON, NCP) were obtained for the Simons Collaborative Marine Atlas Project (CMAP; (58)) with data links provided in the Supplemental Methods. Amplicon samples were paired with corresponding CMAP data as described therein.

### Multi-level Mixed Modeling and Network Analysis

Multivariate linear mixed modeling (MLMM) was performed within the *sommer* package v4.3.3 (59). Weighted gene co-expression network analysis (WGCNA) v1.72-5 (60) was applied to persistent ASVs from eukaryote-containing taxa and Box-Cox–transformed NCP, POC, and PON values to identify ASV modules correlated to each biochemical variable. Details can be found in the Supplemental Methods. Sparse Inverse Covariance Estimation for Ecological Association Inference (SpiecEasi) v1.1.0 (61) was used to infer direct associations between specific persistent eukaryotic and prokaryotic ASVs (Dataset S2). Details can be found in the Supplemental Methods.

## Data Availability

Sequencing and environmental data will be deposited to NCBI and the Simons CMAP repository with accession numbers and DOIs will be added upon public release. All analysis scripts are available at: https://github.com/rkeyMicrobe/picoGrads2025

## Author Contributions

RSK, BPD, and EVA conceived the research project. EVA, JPZ, MRG, HF and BPD planned fieldwork sampling design. RSK led data analysis, data interpretation, figure generation, and manuscript writing, with supervision from BPD. SNC and EVA assisted with refinement of research design. MRG collected seawater samples. RLM, HF and MRG prepared samples for ASV sequencing. All authors contributed to reviewing and revising the manuscript.

## Supporting information

Sample metadata (CSV)

Speic-Easi association index matrix (CSV)

## Acknowledgments

We thank the scientific team and crew of the R/V Kaʻimikai-O-Kanaloa (KOK1606; Gradients 1), R/V Marcus G. Langseth (MGL1704; Gradients 2), and R/V Kilo Moana (KM1906; Gradients 3) and the operational staff of the Simons Collaboration on Ocean Processes and Ecology (SCOPE) team. We also thank Bennet Lambert for early assistance with ASV data processing. This work was supported by grants from the Simons Foundation (Awards 823165 and 999397 to BPD; Award 721244 to EVA; Award 724220 to JPZ; Award 426570SP to EVA and JPZ; Award 00012203 to SNC) as part of the SCOPE Program.

## References

1. Falkowski PG, Wilson C. 1992. Phytoplankton productivity in the North Pacific ocean since 1900 and implications for absorption of anthropogenic CO2. Nature 358:741–743.

2. Finkel ZV, Beardall J, Flynn KJ, Quigg A, Rees TAV, Raven JA. 2010. Phytoplankton in a changing world: cell size and elemental stoichiometry. J Plankton Res 32:119–137.

3. Munk W, Riley G. 1952. Absorption of nutrients by aquatic plants. J Mar Res 11.

4. Bar-On YM, Milo R. 2019. The Biomass Composition of the Oceans: A Blueprint of Our Blue Planet. Cell 179:1451–1454.

5. Sieburth JMcN, Smetacek V, Lenz J. 1978. Pelagic ecosystem structure: Heterotrophic compartments of the plankton and their relationship to plankton size fractions. Limnol Oceanogr 23:1256–1263.

6. Worden A. 2006. Picoeukaryote diversity in coastal waters of the Pacific Ocean. Aquat Microb Ecol 43:165–175.

7. Chisholm SW, Olson RJ, Zettler ER, Goericke R, Waterbury JB, Welschmeyer NA. 1988. A novel free-living prochlorophyte abundant in the oceanic euphotic zone. Nature 334:340–343.

8. Waterbury JB, Watson SW, Guillard RRL, Brand LE. 1979. Widespread occurrence of a unicellular, marine, planktonic, cyanobacterium. | EBSCOhost. https://openurl.ebsco.com/contentitem/doi:10.1038%2F277293a0?sid=ebsco:plink:crawler&id=ebsco:doi:10.1038%2F277293a0. Retrieved 14 February 2025.

9. Vaulot D, Eikrem W, Viprey M, Moreau H. 2008. The diversity of small eukaryotic phytoplankton (≤3 μm) in marine ecosystems. FEMS Microbiol Rev 32:795–820.

10. Worden AZ, Nolan JK, Palenik B. 2004. Assessing the dynamics and ecology of marine picophytoplankton: The importance of the eukaryotic component. Limnol Oceanogr 49:168–179.

11. Buitenhuis ET, Li WKW, Vaulot D, Lomas MW, Landry MR, Partensky F, Karl DM, Ulloa O, Campbell L, Jacquet S, Lantoine F, Chavez F, Macias D, Gosselin M, McManus GB. 2012. Picophytoplankton biomass distribution in the global ocean.

12. Raven JA. 1998. The twelfth Tansley Lecture. Small is beautiful: the picophytoplankton. Funct Ecol 12:503–513.

13. Letscher RT, Moore JK, Martiny AC, Lomas MW. 2023. Biodiversity and Stoichiometric Plasticity Increase Pico-Phytoplankton Contributions to Marine Net Primary Productivity and the Biological Pump. Glob Biogeochem Cycles 37:e2023GB007756.

14. Laws EA, Falkowski PG, Smith Jr. WO, Ducklow H, McCarthy JJ. 2000. Temperature effects on export production in the open ocean. Glob Biogeochem Cycles 14:1231–1246.

15. Follows MJ, Dutkiewicz S, Grant S, Chisholm SW. 2007. Emergent Biogeography of Microbial Communities in a Model Ocean. Science 315:1843–1846.

16. Polovina JJ, Howell EA, Kobayashi DR, Seki MP. 2017. The Transition Zone Chlorophyll Front updated: Advances from a decade of research. Prog Oceanogr 150:79–85.

17. Church MJ, Björkman KM, Karl DM, Saito MA, Zehr JP. 2008. Regional distributions of nitrogen-fixing bacteria in the Pacific Ocean. Limnol Oceanogr 53:63–77.

18. Juranek LW, White AE, Dugenne M, Henderikx Freitas F, Dutkiewicz S, Ribalet F, Ferrón S, Armbrust EV, Karl DM. 2020. The Importance of the Phytoplankton “Middle Class” to Ocean Net Community Production. Glob Biogeochem Cycles 34:e2020GB006702.

19. Kavanaugh MT, Emerson SR, Hales B, Lockwood DM, Quay PD, Letelier RM. 2014. Physicochemical and biological controls on primary and net community production across northeast Pacific seascapes. Limnol Oceanogr 59:2013–2027.

20. Ribalet F, Marchetti A, Hubbard KA, Brown K, Durkin CA, Morales R, Robert M, Swalwell JE, Tortell PD, Armbrust EV. 2010. Unveiling a phytoplankton hotspot at a narrow boundary between coastal and offshore waters. Proc Natl Acad Sci 107:16571–16576.

21. Gradoville MR, Farnelid H, White AE, Turk-Kubo KA, Stewart B, Ribalet F, Ferrón S, Pinedo-Gonzalez P, Armbrust EV, Karl DM, John S, Zehr JP. 2020. Latitudinal constraints on the abundance and activity of the cyanobacterium UCYN-A and other marine diazotrophs in the North Pacific. Limnol Oceanogr 65:1858–1875.

22. Huber P, De Angelis D, Sarmento H, Metz S, Giner CR, Vargas CD, Maiorano L, Massana R, Logares R. 2024. Global distribution, diversity, and ecological niche of Picozoa, a widespread and enigmatic marine protist lineage. Microbiome 12:162.

23. Coale TH, Loconte V, Turk-Kubo KA, Vanslembrouck B, Mak WKE, Cheung S, Ekman A, Chen J-H, Hagino K, Takano Y, Nishimura T, Adachi M, Le Gros M, Larabell C, Zehr JP. 2024. Nitrogen-fixing organelle in a marine alga. Science 384:217–222.

24. Hagino K, Onuma R, Kawachi M, Horiguchi T. 2013. Discovery of an Endosymbiotic Nitrogen-Fixing Cyanobacterium UCYN-A in Braarudosphaera bigelowii (Prymnesiophyceae). PLoS ONE 8:e81749.

25. Zehr JP, Shilova IN, Farnelid HM, Muñoz-Marín MDC, Turk-Kubo KA. 2016. Unusual marine unicellular symbiosis with the nitrogen-fixing cyanobacterium UCYN-A. Nat Microbiol 2:16214.

26. Liefer JD, White AE, Finkel ZV, Irwin AJ, Dugenne M, Inomura K, Ribalet F, Armbrust EV, Karl DM, Fyfe MH, Brown CM, Follows MJ. 2024. Latitudinal patterns in ocean C:N:P reflect phytoplankton acclimation and macromolecular composition. Proc Natl Acad Sci 121:e2404460121.

27. Redfield AC. 1958. The Biological Control of Chemical Factors in the Environment. Am Sci 46:230A–221.

28. Martiny AC, Pham CTA, Primeau FW, Vrugt JA, Moore JK, Levin SA, Lomas MW. 2013. Strong latitudinal patterns in the elemental ratios of marine plankton and organic matter. Nat Geosci 6:279–283.

29. Geider R, and La Roche J. 2002. Redfield revisited: variability of C:N:P in marine microalgae and its biochemical basis. Eur J Phycol 37:1–17.

30. González-García C, Forja J, González-Cabrera MC, Jiménez MP, Lubián LM. 2018. Annual variations of total and fractionated chlorophyll and phytoplankton groups in the Gulf of Cadiz. Sci Total Environ 613–614:1551–1565.

31. Quigg A, Finkel ZV, Irwin AJ, Rosenthal Y, Ho T-Y, Reinfelder JR, Schofield O, Morel FMM, Falkowski PG. 2003. The evolutionary inheritance of elemental stoichiometry in marine phytoplankton. Nature 425:291–294.

32. Ebenezer V, Medlin LK, Ki J-S. 2012. Molecular Detection, Quantification, and Diversity Evaluation of Microalgae. Mar Biotechnol 14:129–142.

33. Guillou L, Eikrem W, Chrétiennot-Dinet M-J, Le Gall F, Massana R, Romari K, Pedrós-Alió C, Vaulot D. 2004. Diversity of Picoplanktonic Prasinophytes Assessed by Direct Nuclear SSU rDNA Sequencing of Environmental Samples and Novel Isolates Retrieved from Oceanic and Coastal Marine Ecosystems. Protist 155:193–214.

34. Romari K, Vaulot D. 2004. Composition and temporal variability of picoeukaryote communities at a coastal site of the English Channel from 18S rDNA sequences. Limnol Oceanogr 49:784–798.

35. Choi DH, An SM, Chun S, Yang EC, Selph KE, Lee CM, Noh JH. 2016. Dynamic changes in the composition of photosynthetic picoeukaryotes in the northwestern Pacific Ocean revealed by high-throughput tag sequencing of plastid 16S rRNA genes. FEMS Microbiol Ecol 92:fiv170.

36. Joli N, Monier A, Logares R, Lovejoy C. 2017. Seasonal patterns in Arctic prasinophytes and inferred ecology of Bathycoccus unveiled in an Arctic winter metagenome. ISME J 11:1372–1385.

37. Lopes dos Santos A, Pollina T, Gourvil P, Corre E, Marie D, Garrido JL, Rodríguez F, Noël M-H, Vaulot D, Eikrem W. 2017. Chloropicophyceae, a new class of picophytoplanktonic prasinophytes. Sci Rep 7:14019.

38. Schön ME, Zlatogursky VV, Singh RP, Poirier C, Wilken S, Mathur V, Strassert JFH, Pinhassi J, Worden AZ, Keeling PJ, Ettema TJG, Wideman JG, Burki F. 2021. Single cell genomics reveals plastid-lacking Picozoa are close relatives of red algae. Nat Commun 12:6651.

39. Brown MW, Heiss AA, Kamikawa R, Inagaki Y, Yabuki A, Tice AK, Shiratori T, Ishida K-I, Hashimoto T, Simpson AGB, Roger AJ. 2018. Phylogenomics Places Orphan Protistan Lineages in a Novel Eukaryotic Super-Group. Genome Biol Evol 10:427–433.

40. Weitz JS, Stock CA, Wilhelm SW, Bourouiba L, Coleman ML, Buchan A, Follows MJ, Fuhrman JA, Jover LF, Lennon JT, Middelboe M, Sonderegger DL, Suttle CA, Taylor BP, Frede Thingstad T, Wilson WH, Eric Wommack K. 2015. A multitrophic model to quantify the effects of marine viruses on microbial food webs and ecosystem processes. ISME J 9:1352–1364.

41. Bachy C, Yung CCM, Needham DM, Gazitúa MC, Roux S, Limardo AJ, Choi CJ, Jorgens DM, Sullivan MB, Worden AZ. 2021. Viruses infecting a warm water picoeukaryote shed light on spatial co-occurrence dynamics of marine viruses and their hosts. ISME J 15:3129–3147.

42. Mayer JA, Taylor FJR. 1979. A virus which lyses the marine nanoflagellate Micromonas pusilla. Nature 281:299–301.

43. Waters RE, Chan AT. 1982. Micromonas pusilla Virus: the Virus Growth Cycle and Associated Physiological Events Within the Host Cells; Host Range Mutation. J Gen Virol 63:199–206.

44. Ban H, Sato S, Yoshikawa S, Yamada K, Nakamura Y, Ichinomiya M, Sato N, Blanc-Mathieu R, Endo H, Kuwata A, Ogata H. 2023. Genome analysis of Parmales, the sister group of diatoms, reveals the evolutionary specialization of diatoms from phago-mixotrophs to photoautotrophs. Commun Biol 6:1–14.

45. Lin Y-C, Campbell T, Chung C-C, Gong G-C, Chiang K-P, Worden AZ. 2012. Distribution Patterns and Phylogeny of Marine Stramenopiles in the North Pacific Ocean. Appl Environ Microbiol 78:3387–3399.

46. Barberán A, Bates ST, Casamayor EO, Fierer N. 2012. Using network analysis to explore co-occurrence patterns in soil microbial communities. ISME J 6:343–351.

47. Horner-Devine MC, Silver JM, Leibold MA, Bohannan BJM, Colwell RK, Fuhrman JA, Green JL, Kuske CR, Martiny JBH, Muyzer G, Øvreås L, Reysenbach A-L, Smith VH. 2007. A Comparison of Taxon Co-Occurrence Patterns for Macro- and Microorganisms. Ecology 88:1345–1353.

48. Worrich A, Musat N, Harms H. 2019. Associational effects in the microbial neighborhood. ISME J 13:2143–2149.

49. Thompson AW, Foster RA, Krupke A, Carter BJ, Musat N, Vaulot D, Kuypers MMM, Zehr JP. 2012. Unicellular Cyanobacterium Symbiotic with a Single-Celled Eukaryotic Alga. Science 337:1546–1550.

50. Moisander PH, Beinart RA, Hewson I, White AE, Johnson KS, Carlson CA, Montoya JP, Zehr JP. 2010. Unicellular Cyanobacterial Distributions Broaden the Oceanic N2 Fixation Domain. Science 327:1512–1514.

51. Caporaso JG, Lauber CL, Walters WA, Berg-Lyons D, Huntley J, Fierer N, Owens SM, Betley J, Fraser L, Bauer M, Gormley N, Gilbert JA, Smith G, Knight R. 2012. Ultra-high-throughput microbial community analysis on the Illumina HiSeq and MiSeq platforms. ISME J 6:1621–1624.

52. Piredda R, Claverie J-M, Decelle J, de Vargas C, Dunthorn M, Edvardsen B, Eikrem W, Forster D, Kooistra WHCF, Logares R, Massana R, Montresor M, Not F, Ogata H, Pawlowski J, Romac S, Sarno D, Stoeck T, Zingone A. 2018. Diatom diversity through HTS-metabarcoding in coastal European seas. Sci Rep 8:18059.

53. Bolyen E, Rideout JR, Dillon MR, Bokulich NA, Abnet CC, Al-Ghalith GA, Alexander H, Alm EJ, Arumugam M, Asnicar F, Bai Y, Bisanz JE, Bittinger K, Brejnrod A, Brislawn CJ, Brown CT, Callahan BJ, Caraballo-Rodríguez AM, Chase J, Cope EK, Da Silva R, Diener C, Dorrestein PC, Douglas GM, Durall DM, Duvallet C, Edwardson CF, Ernst M, Estaki M, Fouquier J, Gauglitz JM, Gibbons SM, Gibson DL, Gonzalez A, Gorlick K, Guo J, Hillmann B, Holmes S, Holste H, Huttenhower C, Huttley GA, Janssen S, Jarmusch AK, Jiang L, Kaehler BD, Kang KB, Keefe CR, Keim P, Kelley ST, Knights D, Koester I, Kosciolek T, Kreps J, Langille MGI, Lee J, Ley R, Liu Y-X, Loftfield E, Lozupone C, Maher M, Marotz C, Martin BD, McDonald D, McIver LJ, Melnik AV, Metcalf JL, Morgan SC, Morton JT, Naimey AT, Navas-Molina JA, Nothias LF, Orchanian SB, Pearson T, Peoples SL, Petras D, Preuss ML, Pruesse E, Rasmussen LB, Rivers A, Robeson MS, Rosenthal P, Segata N, Shaffer M, Shiffer A, Sinha R, Song SJ, Spear JR, Swafford AD, Thompson LR, Torres PJ, Trinh P, Tripathi A, Turnbaugh PJ, Ul-Hasan S, van der Hooft JJJ, Vargas F, Vázquez-Baeza Y, Vogtmann E, von Hippel M, Walters W, Wan Y, Wang M, Warren J, Weber KC, Williamson CHD, Willis AD, Xu ZZ, Zaneveld JR, Zhang Y, Zhu Q, Knight R, Caporaso JG. 2019. Author Correction: Reproducible, interactive, scalable and extensible microbiome data science using QIIME 2. Nat Biotechnol 37:1091–1091.

54. Quast C, Pruesse E, Yilmaz P, Gerken J, Schweer T, Yarza P, Peplies J, Glöckner FO. 2013. The SILVA ribosomal RNA gene database project: improved data processing and web-based tools. Nucleic Acids Res 41:D590–D596.

55. Guillou L, Bachar D, Audic S, Bass D, Berney C, Bittner L, Boutte C, Burgaud G, de Vargas C, Decelle J, del Campo J, Dolan JR, Dunthorn M, Edvardsen B, Holzmann M, Kooistra WHCF, Lara E, Le Bescot N, Logares R, Mahé F, Massana R, Montresor M, Morard R, Not F, Pawlowski J, Probert I, Sauvadet A-L, Siano R, Stoeck T, Vaulot D, Zimmermann P, Christen R. 2013. The Protist Ribosomal Reference database (PR2): a catalog of unicellular eukaryote Small Sub-Unit rRNA sequences with curated taxonomy. Nucleic Acids Res 41:D597–D604.

56. Schloss PD, Westcott SL, Ryabin T, Hall JR, Hartmann M, Hollister EB, Lesniewski RA, Oakley BB, Parks DH, Robinson CJ, Sahl JW, Stres B, Thallinger GG, Van Horn DJ, Weber CF. 2009. Introducing mothur: Open-Source, Platform-Independent, Community-Supported Software for Describing and Comparing Microbial Communities. Appl Environ Microbiol 75:7537–7541.

57. Oksanen J, Simpson GL, Blanchet FG, Kindt R, Legendre P, Minchin PR, O’Hara RB, Solymos P, Stevens MHH, Szoecs E, Wagner H, Barbour M, Bedward M, Bolker B, Borcard D, Carvalho G, Chirico M, Caceres MD, Durand S, Evangelista HBA, FitzJohn R, Friendly M, Furneaux B, Hannigan G, Hill MO, Lahti L, McGlinn D, Ouellette M-H, Cunha ER, Smith T, Stier A, Braak CJFT, Weedon J, Borman T. 2025. vegan: Community Ecology Package (2.6-10).

58. Ashkezari MD, Hagen NR, Denholtz M, Neang A, Burns TC, Morales RL, Lee CP, Hill CN, Armbrust EV. 2021. Simons Collaborative Marine Atlas Project (Simons CMAP): An open-source portal to share, visualize, and analyze ocean data. Limnol Oceanogr Methods 19:488–496.

59. Covarrubias-Pazaran G. 2016. Genome-Assisted Prediction of Quantitative Traits Using the R Package sommer. PLOS ONE 11:e0156744.

60. Langfelder P, Horvath S. 2008. WGCNA: an R package for weighted correlation network analysis. BMC Bioinformatics 9:559.

61. Kurtz ZD, Müller CL, Miraldi ER, Littman DR, Blaser MJ, Bonneau RA. 2015. Sparse and Compositionally Robust Inference of Microbial Ecological Networks. PLOS Comput Biol 11:e1004226.

