## Supplemental Figures, Tables, and Methods for "Picophytoplankton Implicated in Productivity and Biogeochemistry in the North Pacific Transition Zone"

**SUPPLEMENT FIGURES**

S1: Sample Site Depth Profiles for Each Cruise
S2: Eukaryotic 18S Spatial and Temporal Patterns across Yearly Transects
S3: Prokaryotic 16S Spatial and Temporal Patterns across Yearly Transects
S4: Relative Percent Abundance of Major Taxonomic Groups Across Filter Sizes and Regions
S5: WGCNA soft-threshold power and module clustering of persistent phytoplankton Biochemical Variables.
S6: Spiec-Easi-Derived Weight Range and Cluster Assignments for 1° Spiec-Easi Neighbors of Purple and Yellow Candidates

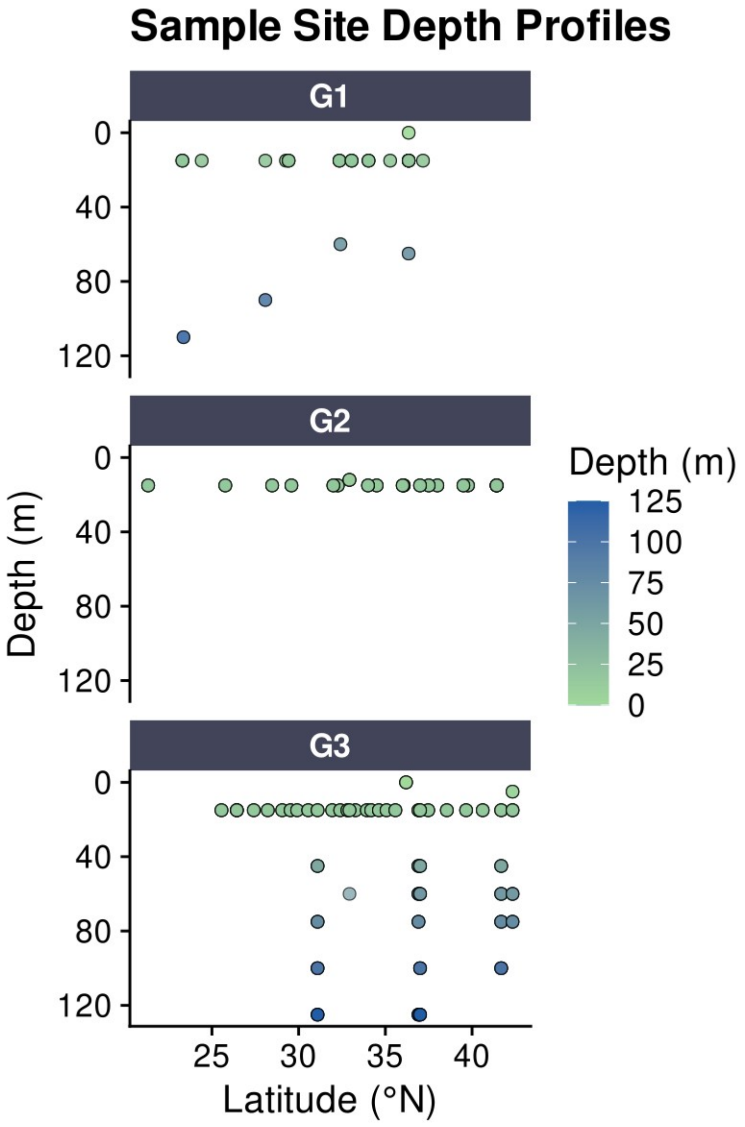


**Fig. S1. Sample Site Depth Profiles for Each Cruise**
Sampling depths (y-axis) and latitudes (x-axis) of all amplicon samples collected during Gradients 1 (G1), Gradients 2 (G2), and Gradients 3 (G3) cruises. Each point represents a unique sampling site that includes both size fractions (0.2–3 µm and >3 µm). Points are colored by collection depth, with light green indicating surface waters and dark blue indicating deeper samples. Samples were collected using either CTD casts or the ship's surface-intake systems. See **Dataset S1** for sample metadata.

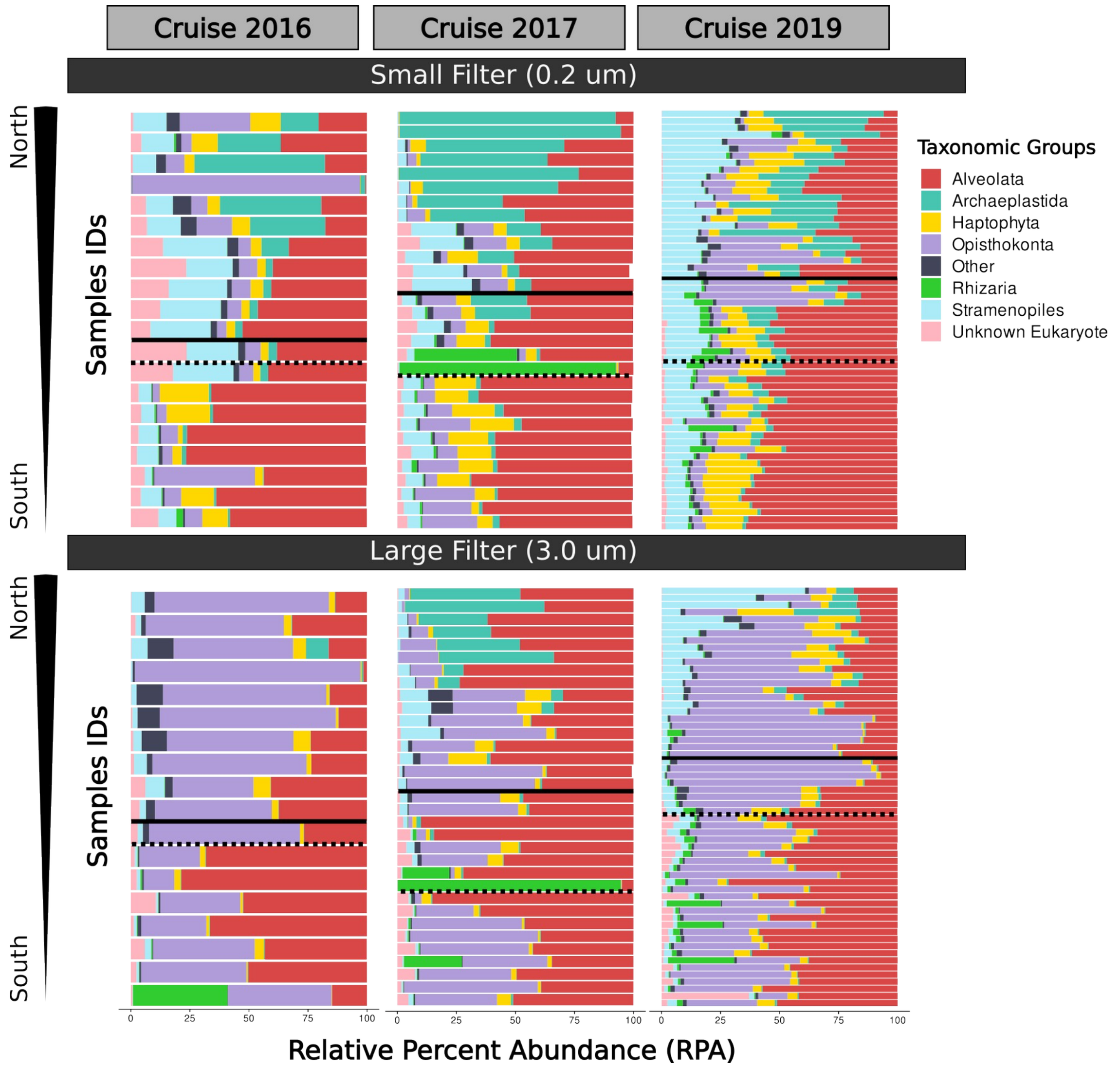


**Fig. S2. Eukaryotic 18S Spatial and Temporal Patterns across Yearly Transects.** Relative percent abundance bar charts for small size fraction samples (top) and large size fraction samples (bottom). Samples are arranged by latitude on the y-axis, and the x-axis displays the total relative percent abundance of each major taxonomic group at the phylum level. 'Unknown Eukaryote' represents eukaryotic ASVs with unknown taxonomic classification at the phylum level. The 'Other' category includes taxonomic groups representing less than 1% of total relative percent abundance per cruise year. The salinity front is denoted by a dotted black line, while the chlorophyll front is denoted by a solid black line.

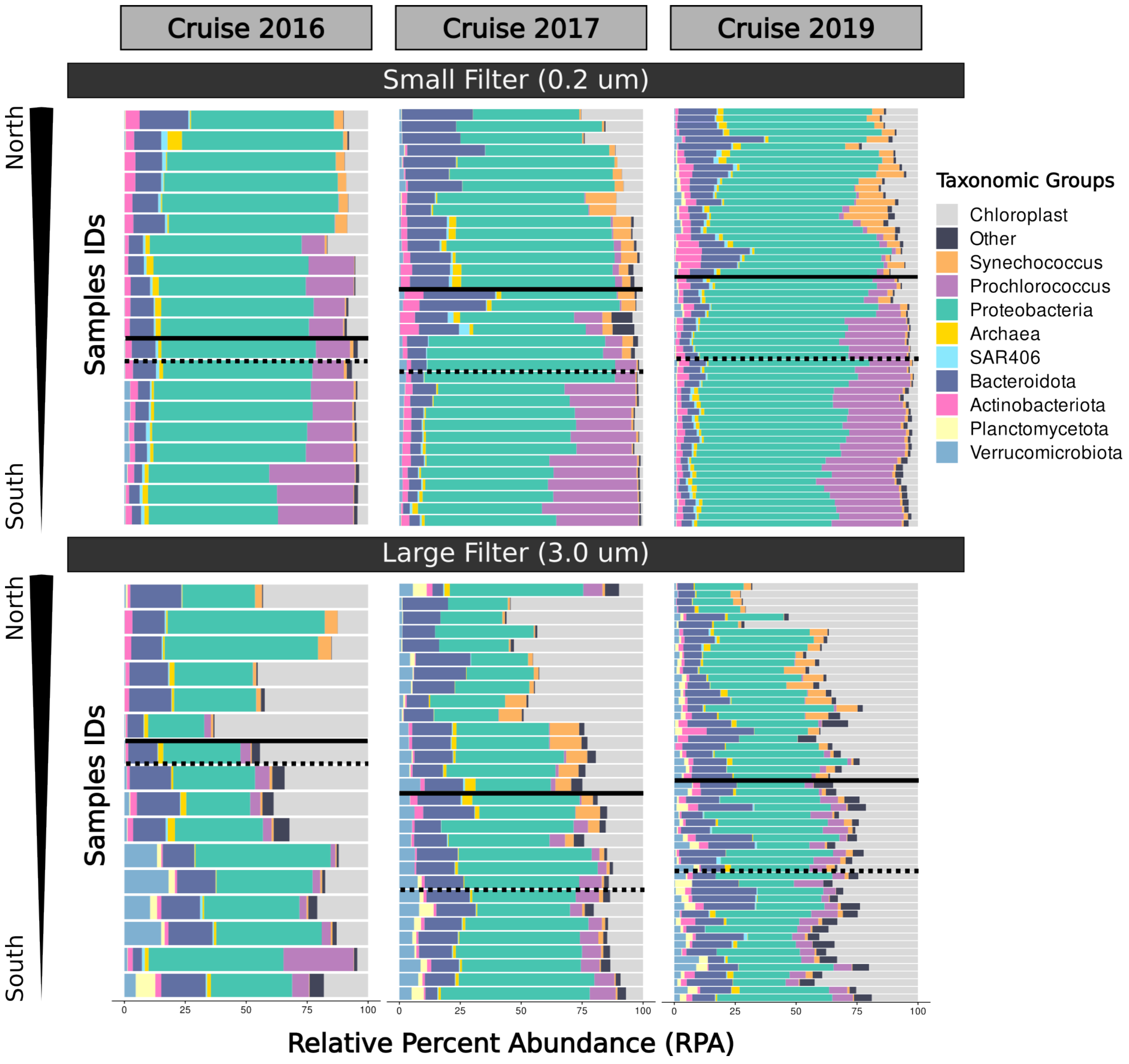


**Fig. S3. Prokaryotic 16S Spatial and Temporal Patterns across Yearly Transects.** Relative percent abundance bar charts for small size fraction samples (top) and large size fraction samples (bottom). Samples are arranged by latitude on the y-axis. The x-axis shows total relative percent abundance of each major taxonomic group at the phylum level. The ‘Prochlorococcus’ and ‘Synechococcus’ categories represent ASVs classified under the respective genera *Prochlorococcus* and *Synechococcus*. The 'Archaea' category groups all ASVs with the domain classification Archaea. The 'Chloroplast' category represents prokaryotic ASVs classified as Chloroplast at the class level. The "Other" category encompasses taxonomic groups representing less than 1% of total relative percent abundance per cruise year, including other cyanobacteria not classified within the *Prochlorococcus* and *Synechococcus* categories. The salinity front is denoted by a dotted black line, while the chlorophyll front is denoted by a solid black line.

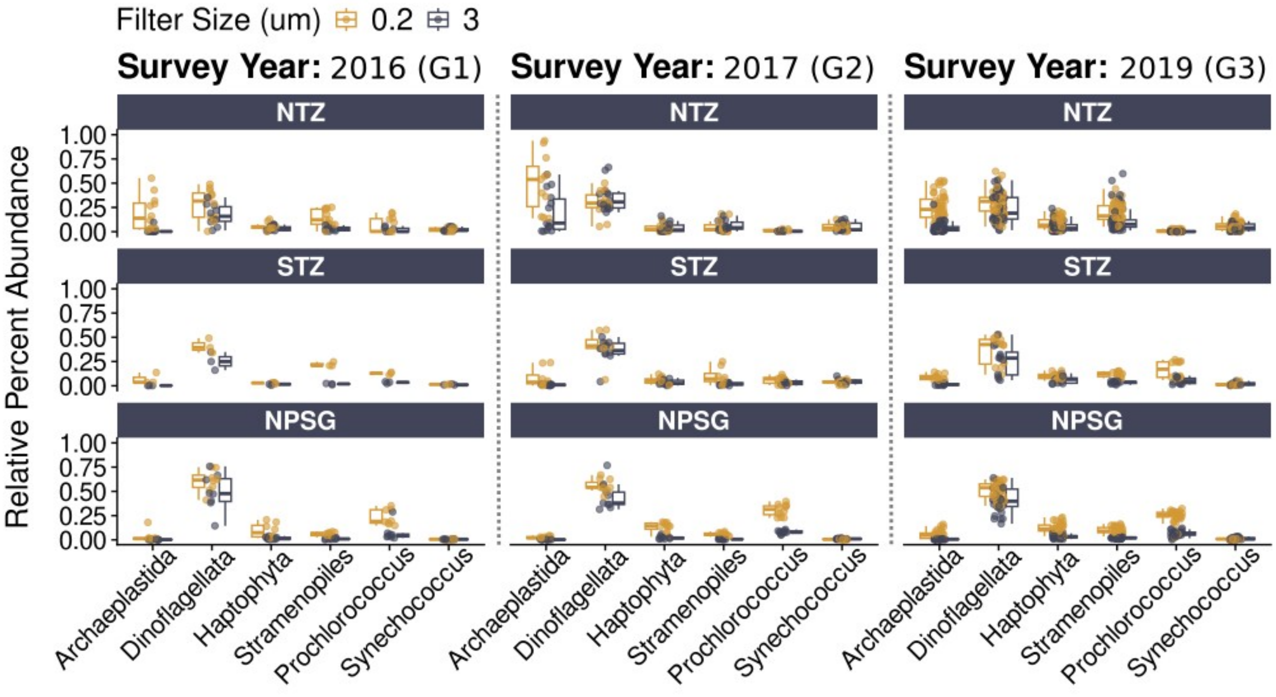


**Fig. S4. Relative Percent Abundance of Major Taxonomic Groups Across Filter Sizes and Regions.** Relative percent abundance of major phytoplankton-containing groups across three regions: NTZ (Northern Transition Zone), STZ (Southern Transition Zone), and NPSG (North Pacific Subtropical Gyre) during three survey years (G1 2016; G2 2017; G3 2019). Values for prokaryotic (*Prochlorococcus* and *Synechococcus*) and eukaryotic (Archaeplastida, Dinoflagellata, Haptophyta, Stramenopiles) relative percent abundances were calculated relative to their corresponding community. Boxplots show the mean and spread of relative percent abundance across samples, with individual samples points plotted adjacent to their respective box plot. Filter size fractions are indicated by gold (0.2 µm) and gray (3 µm). Columns represent each region across the three survey years. Samples from all depths were considered.

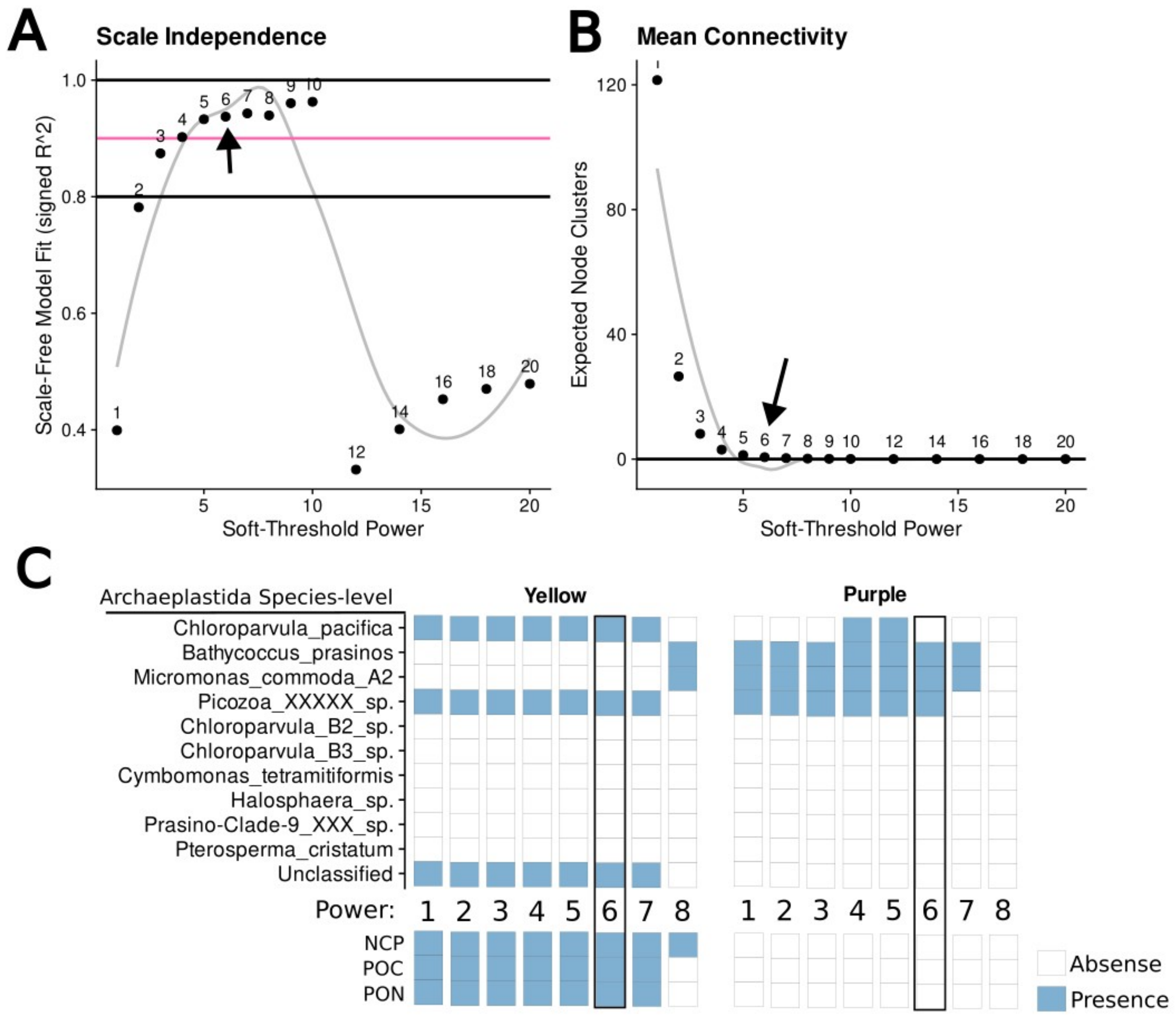


**Fig. S5. WGCNA soft-threshold power and module clustering of persistent phytoplankton Biochemical Variables. (A)** A scale independence plot showing the fit of the network to a scale-free topology across a range of soft-thresholding powers. The y-axis shows the signed $R^2$, which indicates the degree to which the network fits a scale-free topology. A power of 6 was chosen as it provides a high $R^2$ value (above 0.9 represented by the pink line). The dotted lines denote 0.8 and 1.0 $R^2$ values. **(B)** Mean connectivity plot showing the number of expected node clusters, or modules for the different soft-threshold powers. A power of 6 was selected (arrow) as it provides a good balance between high model fit found in Panel A and low mean connectivity that is over zero. **(C)** Heatmap of Archaeplastida species-level ASVs showing their presence (blue) or absence (white) in the clusters associated with biochemical variables (POC, PON, and NCP) across WGCNA soft-thresholding powers ranging from 1 to 8. If the power is too high, the network becomes too sparse, making it harder to detect meaningful connections. If it is too low, weak or random correlations may be kept, making the patterns less reliable. The chosen power reflects a balance between filtering noise and preserving biologically meaningful patterns.

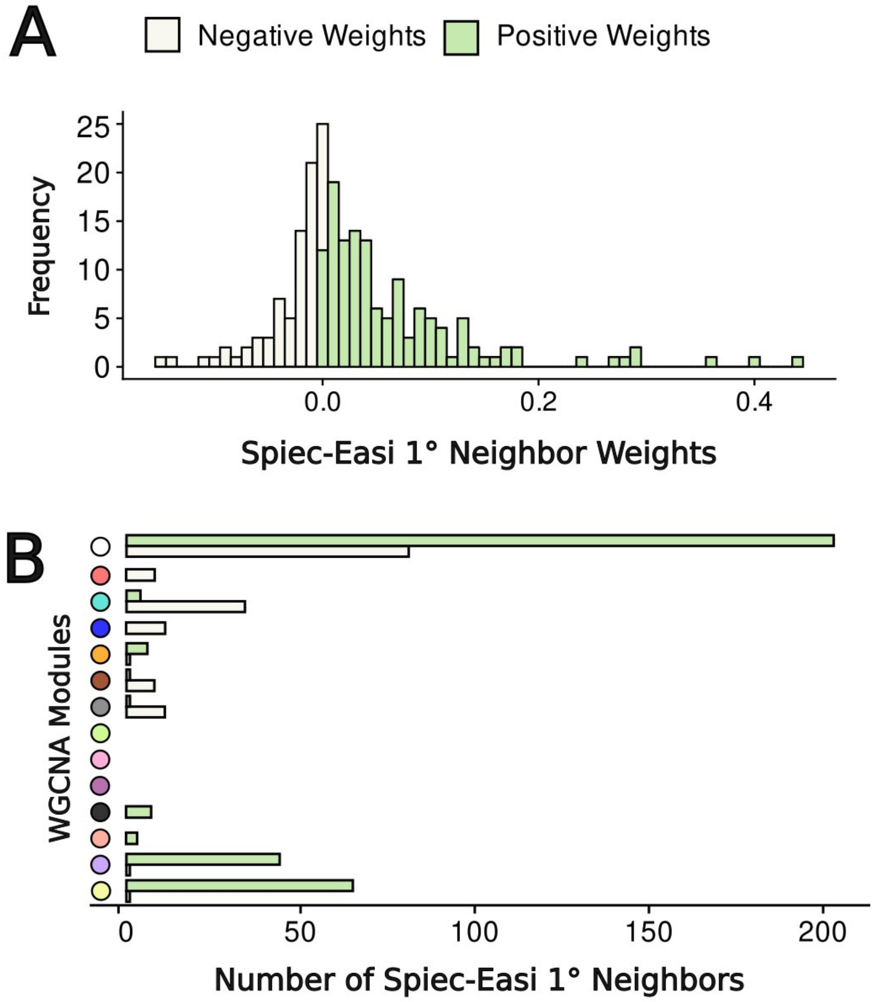


**Fig. S6: Spiec-Easi-Derived Weight Range and Cluster Assignments for 1° Spiec-Easi Neighbors of Purple and Yellow Candidates**
**(A)** Distribution of Spiec-Easi first-degree neighbor weights for ASVs (i.e. Candidates) in the yellow and purple WGCNA clusters. White bars indicate negative weights, while green bars represent positive weights. **(B)** Spiec-Easi first-degree neighbors for candidates found within the yellow and purple WGCNA clusters grouped based on neighbor's WGCNA cluster assignments. The same bar color theme as Panel A is applied. The 'No Assignment' cluster (white) represents ASVs excluded from WGCNA network construction due to quality control filtering for sparsity and heterotrophic bacteria.

**SUPPLEMENT TABLES**

S1:  EnvFit NMDS Categorical Variable Correlations Across Yearly Surveys.
S2:  EnvFit NMDS Correlations on Group Relative Percent Abundance
S3:  Taxonomic Composition of Yellow and Purple WGCNA Clusters
S4: Top 5 Out-degree ASVs for Prokaryotes (16S) and Eukaryotes (18S)
S5:  Yellow and Purple Candidate Connections from Spiec-Easi Network
S6: Rarefaction Thresholds for 18S and 16S Amplicon Sequencing Across the Yearly Surveys

**Table S1. EnvFit NMDS Categorical Variable Correlations Across Yearly Surveys.** Results from an EnvFit analysis performed on eukaryotic and prokaryotic communities across the three yearly surveys (G1 = Cruise 2016, G2 = Cruise 2017, G3 = Cruise 2019). Both $r^2$ values and p-values for 'Filter', 'Region', and 'Depth' variables were evaluated. Higher $r^2$ values indicate stronger correlations between the NMDS ordination and variable. Survey observations are colored to reflect the chosen color palette used in other figures. Cruise 2016 is in green, 2017 in orange, and 2019 in purple.

### EnvFit NMDS Correlations
### Categorical Variables

#### Prokaryotes

| Survey | Factor | r2 | p_value |
|---|---|---|---|
| G1 | Filter | 0.1235 | 0.009 |
| G1 | Region | 0.5019 | 0.001 |
| G1 | Depth | 0.4137 | 0.033 |
| G2 | Filter | 0.0406 | 0.098 |
| G2 | Region | 0.7589 | 0.001 |
| G2 | Depth | 0.0038 | 0.767 |
| G3 | Filter | 0.0936 | 0.001 |
| G3 | Region | 0.4639 | 0.001 |
| G3 | Depth | 0.1808 | 0.001 |

#### Eukaryotes

| Survey | Factor | r2 | p_value |
|---|---|---|---|
| G1 | Filter | 0.2823 | 0.001 |
| G1 | Region | 0.4678 | 0.001 |
| G1 | Depth | 0.0712 | 0.843 |
| G2 | Filter | 0.1684 | 0.001 |
| G2 | Region | 0.3832 | 0.001 |
| G2 | Depth | 0.0800 | 0.007 |
| G3 | Filter | 0.3723 | 0.001 |
| G3 | Region | 0.4042 | 0.001 |
| G3 | Depth | 0.1324 | 0.001 |

**Table S2. EnvFit NMDS Correlations on Group Relative Percent Abundance.**
Results from an EnvFit analysis which was applied to major taxonomic groups of eukaryotic and prokaryotic communities across the three yearly surveys (G1 = Cruise 2016, G2 = Cruise 2017, G3 = Cruise 2019). Other details are the same as Supplement Table 1.0. Details on taxonomic groupings can be found in Fig S2 and 3.

EnvFit NMDS Correlations
Taxa (Continuous) Variables

Prokaryotes

| Survey | Taxa | r2 | p_value |
|---|---|---|---|
| G1 | Actinobacteriota | 0.4017 | 0.001 |
| G1 | Archaea | 0.0100 | 0.840 |
| G1 | Bacteroidota | 0.3729 | 0.002 |
| G1 | Other | 0.3214 | 0.005 |
| G1 | Planctomycetota | 0.3689 | 0.001 |
| G1 | Prochlorococcus | 0.5345 | 0.001 |
| G1 | Proteobacteria | 0.4819 | 0.002 |
| G1 | SAR406 | 0.4289 | 0.001 |
| G1 | Synechococcus | 0.4469 | 0.001 |
| G1 | Verrucomicrobiota | 0.6447 | 0.001 |
| G2 | Actinobacteriota | 0.4473 | 0.001 |
| G2 | Archaea | 0.3344 | 0.001 |
| G2 | Bacteroidota | 0.4571 | 0.001 |
| G2 | Other | 0.0675 | 0.121 |
| G2 | Planctomycetota | 0.3859 | 0.001 |
| G2 | Prochlorococcus | 0.5557 | 0.001 |
| G2 | Proteobacteria | 0.2218 | 0.001 |
| G2 | SAR406 | 0.2987 | 0.001 |
| G2 | Synechococcus | 0.4298 | 0.001 |
| G2 | Verrucomicrobiota | 0.1903 | 0.003 |
| G3 | Actinobacteriota | 0.0316 | 0.176 |
| G3 | Archaea | 0.1512 | 0.001 |
| G3 | Bacteroidota | 0.2533 | 0.001 |
| G3 | Other | 0.3419 | 0.001 |
| G3 | Planctomycetota | 0.1436 | 0.001 |
| G3 | Prochlorococcus | 0.2730 | 0.001 |
| G3 | Proteobacteria | 0.1258 | 0.001 |
| G3 | SAR406 | 0.0804 | 0.005 |
| G3 | Synechococcus | 0.2287 | 0.001 |
| G3 | Verrucomicrobiota | 0.1074 | 0.007 |

Eukaryotes

| Survey | Taxa | r2 | p_value |
|---|---|---|---|
| G1 | Alveolata | 0.7638 | 0.001 |
| G1 | Archaeplastida | 0.3428 | 0.001 |
| G1 | Haptophyta | 0.3312 | 0.001 |
| G1 | Opisthokonta | 0.8372 | 0.001 |
| G1 | Other | 0.2943 | 0.004 |
| G1 | Rhizaria | 0.1834 | 0.019 |
| G1 | Stramenopiles | 0.5232 | 0.001 |
| G1 | Unknown Eukaryote | 0.2900 | 0.004 |
| G2 | Alveolata | 0.2596 | 0.001 |
| G2 | Archaeplastida | 0.7301 | 0.001 |
| G2 | Haptophyta | 0.2257 | 0.001 |
| G2 | Opisthokonta | 0.5000 | 0.001 |
| G2 | Other | 0.0154 | 0.646 |
| G2 | Rhizaria | 0.3012 | 0.001 |
| G2 | Stramenopiles | 0.0709 | 0.100 |
| G2 | Unknown Eukaryote | 0.2856 | 0.001 |
| G3 | Alveolata | 0.7113 | 0.001 |
| G3 | Archaeplastida | 0.5673 | 0.001 |
| G3 | Haptophyta | 0.3922 | 0.001 |
| G3 | Opisthokonta | 0.7304 | 0.001 |
| G3 | Other | 0.1602 | 0.001 |
| G3 | Rhizaria | 0.1035 | 0.003 |
| G3 | Stramenopiles | 0.6061 | 0.001 |
| G3 | Unknown Eukaryote | 0.1947 | 0.001 |

**Table S3. Taxonomic Composition of Yellow and Purple WGCNA Clusters.**
Taxonomy information for ASVs identified as members of the Yellow and Purple clusters the WGCNA network. Each row is an individual ASV where cells in the ‘Cluster’ column are colored based on WGCNA cluster. Biochemical variables NCP, POC, and PON are included in the ‘ASV_ID’ column to show their cluster assignment.

| Cluster | ASV_ID | Group | Phylum | Class | Order | Family | Genus | Species |
|---|---|---|---|---|---|---|---|---|
| Purple | ASV2f69e | Archaeplastida | Chlorophyta | Mamiellophyceae | Mamiellales | Mamiellaceae | Micromonas | Micromonas_commoda_A2 |
| Purple | ASV405f6 | Archaeplastida | Chlorophyta | Mamiellophyceae | Mamiellales | Bathycoccaceae | Bathycoccus | Bathycoccus_prasinos |
| Purple | ASV7c01a | Dinoflagellata | Dinoflagellata | Syndiniales | Dino-Group-II | Dino-Group-II-Clade-10-and-11 | Dino-Group-II-Clade-10-and-11_X | Dino-Group-II-Clade-10-and-11_X_sp. |
| Purple | ASV4963e | Dinoflagellata | Dinoflagellata | Syndiniales | Dino-Group-I | Dino-Group-I-Clade-4 | Dino-Group-I-Clade-4_X | Dino-Group-I-Clade-4_X_sp. |
| Purple | ASV25f1c | Dinoflagellata | Dinoflagellata | Dinophyceae | | | | |
| Purple | ASVf6ee1 | Stramenopiles | Gyrista | Pelagophyceae | Pelagomonadales | Pelagomonadaceae | Pelagomonas | Pelagomonas_calceolata |
| Purple | ASV4d8f7 | Stramenopiles | Gyrista | Bolidophyceae | Parmales | Triparmaceae | Triparma | Triparma_pacifica |
| Purple | ASV0ec07 | Stramenopiles | Gyrista | Pelagophyceae | Pelagomonadales | Pelagomonadaceae | Pelagomonadaceae_clade_C | Pelagomonadaceae_clade_C_sp. |
| Purple | ASVb472b | Stramenopiles | Gyrista | Pelagophyceae | Pelagomonadales | Pelagomonadales_clade_B | Pelagomonadales_clade_B1 | Pelagomonadales_clade_B1_sp. |
| Purple | ASVd5d61 | Stramenopiles | Gyrista | Pelagophyceae | Pelagomonadales | Pelagomonadaceae | Aureococcus | Aureococcus_anophagefferens |
| Purple | ASVe1e58 | Stramenopiles | Gyrista | Mediophyceae | Cymatosirales | Cymatosiraceae | Brockmanniella | Brockmanniella_brockmannii |
| Purple | ASV06b13 | Stramenopiles | Gyrista | Bacillariophyceae | Bacillariales | Bacillariaceae | Fragilariopsis | |
| Purple | ASVec9cb | Stramenopiles | Gyrista | Gyrista_X | Gyrista_XX | MAST-2 | MAST-2D | MAST-2D_sp. |
| Purple | ASVc675c | Stramenopiles | Gyrista | Dictyochophyceae | Dictyochophyceae_X | Pedinellales | | |
| Purple | ASV87490 | Stramenopiles | Gyrista | Dictyochophyceae | Dictyochophyceae_X | Dictyochales | Dictyocha | Dictyocha_speculum |
| Purple | ASVbb0ee | Haptophyta | Haptophyta | Prymnesiophyceae | Prymnesiophyceae_Clade_D | Prymnesiophyceae_Clade_D_X | Prymnesiophyceae_Clade_D_XX | Prymnesiophyceae_Clade_D_XX_sp. |
| Purple | ASV98a19 | Haptophyta | Haptophyta | Prymnesiophyceae | Prymnesiales | Chrysochromulinaceae | Chrysochromulina | Chrysochromulina_sp. |
| Purple | ASV7fd9a | Haptophyta | Haptophyta | Prymnesiophyceae | Phaeocystales | Phaeocystaceae | Phaeocystis | Phaeocystis_pouchetii |
| Purple | ASV94519 | Archaeplastida | Picozoa | Picozoa_XX | Picozoa_XXX | Picozoa_XXXX | Picozoa_XXXXX | Picozoa_XXXXX_sp. |
| Yellow | ASV1c13e | Stramenopiles | Bigyra | Sagenista | Sagenista_X | MAST-7 | MAST-7A | MAST-7A_sp. |
| Yellow | ASVa1d5e | Archaeplastida | Chlorophyta | Chloropicophyceae | Chloropicales | Chloropicaceae | Chloroparvula | Chloroparvula_pacifica |
| Yellow | ASV5f4ba | Synechococcus | Cyanobacteria | Cyanobacteriia | Synechococcales | Cyanobiaceae | Synechococcus_CC9902 | |
| Yellow | ASV62139 | Synechococcus | Cyanobacteria | Cyanobacteriia | Synechococcales | Cyanobiaceae | Synechococcus_CC9902 | uncultured_sp |
| Yellow | ASV6efcd | Synechococcus | Cyanobacteria | Cyanobacteriia | Synechococcales | Cyanobiaceae | Synechococcus_CC9902 | |
| Yellow | ASV3009d | Synechococcus | Cyanobacteria | Cyanobacteriia | Synechococcales | Cyanobiaceae | Synechococcus_CC9902 | uncultured_sp |
| Yellow | ASV4bf35 | Dinoflagellata | Dinoflagellata | Syndiniales | Dino-Group-II | Dino-Group-II-Clade-10-and-11 | Dino-Group-II-Clade-10-and-11_X | Dino-Group-II-Clade-10-and-11_X_sp. |
| Yellow | ASVab7c2 | Dinoflagellata | Dinoflagellata | Dinophyceae | | | | |
| Yellow | ASV69bae | Stramenopiles | Gyrista | Bolidophyceae | Parmales | Parmales_env_3 | Parmales_env_3A | Parmales_env_3A_sp. |
| Yellow | ASV6f81e | Stramenopiles | Gyrista | Dictyochophyceae | Dictyochophyceae_X | Florenciellales | Pseudochattonella | Pseudochattonella_sp. |
| Yellow | ASV74072 | Stramenopiles | Gyrista | Gyrista_X | Gyrista_XX | MAST-1 | MAST-1B | MAST-1B_sp. |
| Yellow | ASVee3a5 | Stramenopiles | Gyrista | Gyrista_X | Gyrista_XX | MAST-1 | MAST-1C | MAST-1C_sp. |
| Yellow | ASVf6b94 | Stramenopiles | Gyrista | Dictyochophyceae | Dictyochophyceae_X | Dictyochophyceae_XX | Dictyochophyceae_XXX | Dictyochophyceae_XXX_sp. |
| Yellow | ASVaff6d | Stramenopiles | Gyrista | Mediophyceae | | | | |
| Yellow | ASV064f4 | Stramenopiles | Gyrista | Pelagophyceae | Pelagomonadales | Pelagomonadales_clade_B | Pelagomonadales_clade_B1 | Pelagomonadales_clade_B1_sp. |
| Yellow | ASVa381f | Stramenopiles | Gyrista | Dictyochophyceae | Dictyochophyceae_X | Florenciellales | Pseudochattonella | Pseudochattonella_sp. |
| Yellow | ASVc0b80 | Stramenopiles | Gyrista | Gyrista_X | Gyrista_XX | MAST-1 | MAST-1A | MAST-1A_sp. |
| Yellow | ASV19672 | Haptophyta | Haptophyta | Prymnesiophyceae | Prymnesiales | Chrysochromulinaceae | Chrysochromulina | Chrysochromulina_sp. |
| Yellow | ASV3814e | Haptophyta | Haptophyta | Haptophyta_Clade_HAP3 | Haptophyta_Clade_HAP3_X | Haptophyta_Clade_HAP3_XX | Haptophyta_Clade_HAP3_XXX | Haptophyta_Clade_HAP3_XXX_sp. |
| Yellow | ASVf391e | Haptophyta | Haptophyta | Prymnesiophyceae | Phaeocystales | Phaeocystaceae | Phaeocystis | Phaeocystis_antarctica |
| Yellow | ASV9edcf | Haptophyta | Haptophyta | Prymnesiophyceae | Prymnesiales | Chrysochromulinaceae | Chrysochromulina | Chrysochromulina_sp. |
| Yellow | ASV23dac | Archaeplastida | Picozoa | Picozoa_XX | Picozoa_XXX | Picozoa_XXXX | Picozoa_XXXXX | Picozoa_XXXXX_sp. |
| Yellow | ASV077d6 | Archaeplastida | Prasinodermophyta | Prasinodermophyceae | Prasinodermales | Prasinodermaceae | Prasinoderma | |
| Yellow | NCP | | | | | | | |
| Yellow | POC | | | | | | | |
| Yellow | PON | | | | | | | |

**Table S4. Top 5 Out-degree ASVs for Prokaryotes (16S) and Eukaryotes (18S)**

ASVs with the highest connectivity are highlighted in blue (16S prokaryotes) and green (18S eukaryotes). Outdegree values represent the number of first-degree connections (network edges) for each ASV in the overall SpiecEasi co-occurrence network. ASVs with high outdegree have the most connections in the network. Taxonomic classifications are provided.

**Top 5 OutDegree ASVs**

Cytoscape Network Analysis Tool

| Group | Domain | ASV ID | Outdegree | Phylum | Class | Order | Family | Genus | Species |
|---|---|---|---|---|---|---|---|---|---|
| Proteobacteria | 16S | ASV7d83e | 60 | Proteobacteria | Alphaproteobacteria | SAR11_clade | Clade_II | Clade_II | NA |
| Bacteroidota | 16S | ASV2c871 | 59 | Bacteroidota | Bacteroidia | Flavobacteriales | Flavobacteriaceae | NS5_marine_group | NA |
| Proteobacteria | 16S | ASVaac3f | 56 | Proteobacteria | Alphaproteobacteria | SAR11_clade | Clade_II | Clade_II | NA |
| Actinobacteria | 16S | ASV74a26 | 55 | Actinobacteriota | Actinobacteria | Corynebacteriales | Mycobacteriaceae | Mycobacterium | NA |
| Proteobacteria | 16S | ASV2c674 | 54 | Proteobacteria | Alphaproteobacteria | SAR11_clade | Clade_I | Clade_Ia | NA |
| Haptophyta | 18S | ASV41683 | 47 | Haptophyta | NA | NA | NA | NA | NA |
| Archaeplastida | 18S | ASVa1d5e | 44 | Chlorophyta | Chloropicophyceae | Chloropicales | Chloropicaceae | Chloroparvula | Chloroparvula_pacifica |
| Dinoflagellata | 18S | ASV00a36 | 42 | Dinoflagellata | Syndiniales | Dino-Group-I | Dino-Group-I-Clade-1 | Dino-Group-I-Clade-1_X | Dino-Group-I-Clade-1_X_sp. |
| Stramenopiles | 18S | ASVb3907 | 41 | Gyrista | Dictyochophyceae | Dictyochophyceae_X | Dictyochales | Dictyocha | Dictyocha_globosa |
| Archaeplastida | 18S | ASV94519 | 41 | Picozoa | Picozoa_XX | Picozoa_XXX | Picozoa_XXXX | Picozoa_XXXXX | Picozoa_XXXXX_sp. |

**Table S5. Yellow and Purple Candidate Connections from Spiec-Easi Network.** Top positive-weight connections between yellow and purple WGCNA candidate ASVs and their first-degree neighbors. ASVs are color-coded by their WGCNA cluster assignments. Taxonomic classifications (Family | Genus | Species) are provided.

| Candidate Cluster | Candidate ASV ID | Candidate Group | Candidate Taxonomy (Family : Species) | Weight | Associate Cluster | Associate ASV ID | Associate Domain | Associate Group | Associate Taxonomy (Family : Species) |
|---|---|---|---|---|---|---|---|---|---|
| Cyan | asv180a1 | Haptophyta | Braarudosphaeraceae \| Braarudosphaeraceae_X \| Braarudosphaeraceae_X_sp. | 0.18 | Brown | asvc701f | Euk | Haptophyta | Braarudosphaeraceae \| Braarudosphaera \| Braarudosphaera_bigelowii |
| Cyan | asv180a1 | Haptophyta | Braarudosphaeraceae \| Braarudosphaeraceae_X \| Braarudosphaeraceae_X_sp. | 0.24 | Unassigned | asv4d1ff | Prok | UCYN_A | Microcystaceae \| Atelocyanobacterium_(UCYN-A) \| Candidatus_Atelocyanobacterium |
| Purple | asv06b13 | Stramenopiles | Bacillariaceae \| Fragilariopsis \| NA | 0.08 | Purple | asv7fd9a | Euk | Haptophyta | Phaeocystaceae \| Phaeocystis \| Phaeocystis_pouchetii |
| Purple | asv405f6 | Archaeplastida | Bathycoccaceae \| Bathycoccus \| Bathycoccus_prasinos | 0.44 | Purple | asvf6ee1 | Euk | Stramenopiles | Pelagomonadaceae \| Pelagomonas \| Pelagomonas_calceolata |
| Purple | asv405f6 | Archaeplastida | Bathycoccaceae \| Bathycoccus \| Bathycoccus_prasinos | 0.13 | Unassigned | asve710b | Prok | Gammaproteobacteria | Spongiibacteraceae \| BD1-7_clade \| uncultured_sp |
| Purple | asv98a19 | Haptophyta | Chrysochromulinaceae \| Chrysochromulina \| Chrysochromulina_sp. | 0.20 | Unassigned | asv705d3 | Euk | Dinoflagellata | NA \| NA \| NA |
| Purple | asve1e58 | Stramenopiles | Cymatosiraceae \| Brockmanniella \| Brockmanniella_brockmannii | 0.10 | Purple | asv87490 | Euk | Stramenopiles | Dictyochales \| Dictyocha \| Dictyocha_speculum |
| Purple | asv87490 | Stramenopiles | Dictyochales \| Dictyocha \| Dictyocha_speculum | 0.06 | Unassigned | asv7ac0a | Euk | Dinoflagellata | NA \| NA \| NA |
| Purple | asv4963e | Dinoflagellata | Dino-Group-I-Clade-4 \| Dino-Group-I-Clade-4_X \| Dino-Group-I-Clade-4_X_sp. | 0.29 | Clay | asv3b1aa | Euk | Archaeplastida | Chloropicaceae \| Chloroparvula \| Chloroparvula_B3_sp. |
| Purple | asv7c01a | Dinoflagellata | Dino-Group-II-Clade-10-and-11 \| Dino-Group-II-Clade-10-and-11_X \| Dino-Group-II-Clade-10-and-11_X_sp. | 0.06 | Unassigned | asv097e8 | Euk | Dinoflagellata | NA \| NA \| NA |
| Purple | asvec9cb | Stramenopiles | MAST-2 \| MAST-2D \| MAST-2D_sp. | 0.07 | Yellow | asv23dac | Euk | Archaeplastida | Picozoa_XXXX \| Picozoa_XXXXX \| Picozoa_XXXXX_sp. |
| Purple | asv2f69e | Archaeplastida | Mamiellaceae \| Micromonas \| Micromonas_commoda_A2 | 0.27 | Purple | asv7fd9a | Euk | Haptophyta | Phaeocystaceae \| Phaeocystis \| Phaeocystis_pouchetii |
| Purple | asv2f69e | Archaeplastida | Mamiellaceae \| Micromonas \| Micromonas_commoda_A2 | 0.13 | Unassigned | asvd167b | Prok | Alphaproteobacteria | SAR116_clade \| Candidatus_Puniceispirillum \| NA |
| Purple | asv25f1c | Dinoflagellata | NA \| NA \| NA | 0.09 | Yellow | asv69bae | Euk | Stramenopiles | Parmales_env_3 \| Parmales_env_3A \| Parmales_env_3A_sp. |
| Purple | asvc675c | Stramenopiles | Pedinellales \| NA \| NA | 0.06 | Purple | asv87490 | Euk | Stramenopiles | Dictyochales \| Dictyocha \| Dictyocha_speculum |
| Purple | asvd5d61 | Stramenopiles | Pelagomonadaceae \| Aureococcus \| Aureococcus_anophagefferens | 0.09 | Purple | asve1e58 | Euk | Stramenopiles | Cymatosiraceae \| Brockmanniella \| Brockmanniella_brockmannii |
| Purple | asv0ec07 | Stramenopiles | Pelagomonadaceae \| Pelagomonadaceae_clade_C \| Pelagomonadaceae_clade_C_sp. | 0.14 | Purple | asvb472b | Euk | Stramenopiles | Pelagomonadales_clade_B \| Pelagomonadales_clade_B1 \| Pelagomonadales_clade_B1_sp. |
| Purple | asvf6ee1 | Stramenopiles | Pelagomonadaceae \| Pelagomonas \| Pelagomonas_calceolata | 0.36 | Purple | asvbb0ee | Euk | Haptophyta | Prymnesiophyceae_Clade_D_X \| Prymnesiophyceae_Clade_D_XX \| Prymnesiophyceae_Clade_D_XX_sp. |
| Purple | asvb472b | Stramenopiles | Pelagomonadales_clade_B \| Pelagomonadales_clade_B1 \| Pelagomonadales_clade_B1_sp. | 0.11 | Purple | asv87490 | Euk | Stramenopiles | Dictyochales \| Dictyocha \| Dictyocha_speculum |
| Purple | asv7fd9a | Haptophyta | Phaeocystaceae \| Phaeocystis \| Phaeocystis_pouchetii | 0.28 | Yellow | asvf391e | Euk | Haptophyta | Phaeocystaceae \| Phaeocystis \| Phaeocystis_antarctica |
| Purple | asv94519 | Archaeplastida | Picozoa_XXXX \| Picozoa_XXXXX \| Picozoa_XXXXX_sp. | 0.11 | Black | asv85495 | Euk | Dinoflagellata | Dino-Group-III_X \| Dino-Group-III_XX \| Dino-Group-III_XX_sp. |
| Purple | asv94519 | Archaeplastida | Picozoa_XXXX \| Picozoa_XXXXX \| Picozoa_XXXXX_sp. | 0.09 | Unassigned | asvc8e36 | Prok | Actinobacteriota | Actinomarinaceae \| Candidatus_Actinomarina \| uncultured_sp |
| Purple | asvbb0ee | Haptophyta | Prymnesiophyceae_Clade_D_X \| Prymnesiophyceae_Clade_D_XX \| Prymnesiophyceae_Clade_D_XX_sp. | 0.19 | Unassigned | asv2ddf7 | Euk | Haptophyta | Chrysochromulinaceae \| Chrysochromulina \| NA |
| Purple | asv4d8f7 | Stramenopiles | Triparmaceae \| Triparma \| Triparma_pacifica | 0.04 | Purple | asv25f1c | Euk | Dinoflagellata | NA \| NA \| NA |
| Yellow | asva1d5e | Archaeplastida | Chloropicaceae \| Chloroparvula \| Chloroparvula_pacifica | 0.37 | Unassigned | asvb10a4 | Euk | Dinoflagellata | Dino-Group-I-Clade-1 \| Dino-Group-I-Clade-1_X \| Dino-Group-I-Clade-1_X_sp. |
| Yellow | asva1d5e | Archaeplastida | Chloropicaceae \| Chloroparvula \| Chloroparvula_pacifica | 0.09 | Unassigned | asv0f593 | Prok | Bacteroidota | Flavobacteriaceae \| NS2b_marine_group \| NA |
| Yellow | asv19672 | Haptophyta | Chrysochromulinaceae \| Chrysochromulina \| Chrysochromulina_sp. | 0.13 | Yellow | asv23dac | Euk | Archaeplastida | Picozoa_XXXX \| Picozoa_XXXXX \| Picozoa_XXXXX_sp. |
| Yellow | asv9edcf | Haptophyta | Chrysochromulinaceae \| Chrysochromulina \| Chrysochromulina_sp. | 0.18 | Yellow | asvc0b80 | Euk | Stramenopiles | MAST-1 \| MAST-1A \| MAST-1A_sp. |
| Yellow | asv5f4ba | Synechococcus | Cyanobiaceae \| Synechococcus_CC9902 \| NA | 0.15 | Unassigned | asv9feeb | Euk | Stramenopiles | Thalassiosiraceae \| NA \| NA |
| Yellow | asv5f4ba | Synechococcus | Cyanobiaceae \| Synechococcus_CC9902 \| NA | 0.12 | Yellow | asv6efcd | Prok | Synechococcus | Cyanobiaceae \| Synechococcus_CC9902 \| NA |
| Yellow | asv6efcd | Synechococcus | Cyanobiaceae \| Synechococcus_CC9902 \| NA | 0.05 | Purple | asve1e58 | Euk | Stramenopiles | Cymatosiraceae \| Brockmanniella \| Brockmanniella_brockmannii |
| Yellow | asv3009d | Synechococcus | Cyanobiaceae \| Synechococcus_CC9902 \| uncultured_sp | 0.20 | Unassigned | asv357e8 | Euk | Stramenopiles | Chrysophyceae_Clade-EC2H_X \| Chrysophyceae_Clade-EC2H_XX \| Chrysophyceae_Clade-EC2H_XX_sp. |
| Yellow | asv3009d | Synechococcus | Cyanobiaceae \| Synechococcus_CC9902 \| uncultured_sp | 0.14 | Unassigned | asv3b154 | Prok | Alphaproteobacteria | Clade_I \| NA \| NA |
| Yellow | asv62139 | Synechococcus | Cyanobiaceae \| Synechococcus_CC9902 \| uncultured_sp | 0.17 | Yellow | asv69bae | Euk | Stramenopiles | Parmales_env_3 \| Parmales_env_3A \| Parmales_env_3A_sp. |
| Yellow | asv62139 | Synechococcus | Cyanobiaceae \| Synechococcus_CC9902 \| uncultured_sp | 0.07 | Unassigned | asv2d102 | Prok | Cyanobacteria | Cyanobiaceae \| NA \| NA |
| Yellow | asvf6b94 | Stramenopiles | Dictyochophyceae_XX \| Dictyochophyceae_XXX \| Dictyochophyceae_XXX_sp. | 0.14 | Unassigned | asv57c31 | Euk | Dinoflagellata | Dino-Group-II-Clade-7 \| Dino-Group-II-Clade-7_X \| Dino-Group-II-Clade-7_X_sp. |
| Yellow | asv4bf35 | Dinoflagellata | Dino-Group-II-Clade-10-and-11 \| Dino-Group-II-Clade-10-and-11_X \| Dino-Group-II-Clade-10-and-11_X_sp. | 0.13 | Yellow | asv69bae | Euk | Stramenopiles | Parmales_env_3 \| Parmales_env_3A \| Parmales_env_3A_sp. |
| Yellow | asv6f81e | Stramenopiles | Florenciellales \| Pseudochattonella \| Pseudochattonella_sp. | 0.07 | Yellow | asva381f | Euk | Stramenopiles | Florenciellales \| Pseudochattonella \| Pseudochattonella_sp. |
| Yellow | asva381f | Stramenopiles | Florenciellales \| Pseudochattonella \| Pseudochattonella_sp. | 0.07 | Unassigned | asv73f97 | Euk | Stramenopiles | Thalassiosiraceae \| Thalassiosira \| NA |
| Yellow | asv3814e | Haptophyta | Haptophyta_Clade_HAP3_XX \| Haptophyta_Clade_HAP3_XXX \| Haptophyta_Clade_HAP3_XXX_sp. | 0.04 | Yellow | asvab7c2 | Euk | Dinoflagellata | NA \| NA \| NA |
| Yellow | asv74072 | Stramenopiles | MAST-1 \| MAST-1B \| MAST-1B_sp. | 0.40 | Yellow | asvee3a5 | Euk | Stramenopiles | MAST-1 \| MAST-1C \| MAST-1C_sp. |
| Yellow | asvee3a5 | Stramenopiles | MAST-1 \| MAST-1C \| MAST-1C_sp. | 0.11 | Yellow | asvf6b94 | Euk | Stramenopiles | Dictyochophyceae_XX \| Dictyochophyceae_XXX \| Dictyochophyceae_XXX_sp. |
| Yellow | asv1c13e | Stramenopiles | MAST-7 \| MAST-7A \| MAST-7A_sp. | 0.05 | Yellow | asvc0b80 | Euk | Stramenopiles | MAST-1 \| MAST-1A \| MAST-1A_sp. |
| Yellow | asvaff6d | Stramenopiles | NA \| NA \| NA | 0.29 | Yellow | asv9edcf | Euk | Haptophyta | Chrysochromulinaceae \| Chrysochromulina \| Chrysochromulina_sp. |
| Yellow | asv69bae | Stramenopiles | Parmales_env_3 \| Parmales_env_3A \| Parmales_env_3A_sp. | 0.11 | Yellow | asvc0b80 | Euk | Stramenopiles | MAST-1 \| MAST-1A \| MAST-1A_sp. |
| Yellow | asvf391e | Haptophyta | Phaeocystaceae \| Phaeocystis \| Phaeocystis_antarctica | 0.03 | Yellow | asv9edcf | Euk | Haptophyta | Chrysochromulinaceae \| Chrysochromulina \| Chrysochromulina_sp. |
| Yellow | asv23dac | Archaeplastida | Picozoa_XXXX \| Picozoa_XXXXX \| Picozoa_XXXXX_sp. | 0.15 | Yellow | asv1c13e | Euk | Stramenopiles | MAST-7 \| MAST-7A \| MAST-7A_sp. |
| Yellow | asv077d6 | Archaeplastida | Prasinodermaceae \| Prasinoderma \| NA | 0.16 | Purple | asv0ec07 | Euk | Stramenopiles | Pelagomonadaceae \| Pelagomonadaceae_clade_C \| Pelagomonadaceae_clade_C_sp. |
| Yellow | asv077d6 | Archaeplastida | Prasinodermaceae \| Prasinoderma \| NA | 0.11 | Unassigned | asv3c25a | Prok | Gammaproteobacteria | Vibrionaceae \| NA \| NA |

**Table S6. Rarefaction Thresholds for 18S and 16S Amplicon Sequencing Across the Yearly Surveys.** The table shows rarefaction thresholds applied to eukaryotic 18S and prokaryotic 16S ASV datasets for three yearly surveys collected during 2016–2019 survey years. Thresholds were selected based on sequencing depth to ensure comparability across samples. Cruise 2016 (G1) is represented in green, Cruise 2017 (G2) in orange, and Cruise 2019 (G3) in purple.

| Cruise | Prokaryotes_16S | Eukaryotes_18S |
|---|---:|---:|
| 2016_Gradients1 | 90,000 | 35,000 |
| 2017_Gradients2 | 20,000 | 10,000 |
| 2019_Gradients3 | 50,000 | 70,000 |

**DATASET FILES**
S1: Sample Site Meta (Replicates)
S2: Speic-Easi Associations

Also available on https://github.com/rkeyMicrobe/picoGrads2025

**SUPPLEMENT MATERIALS AND METHODS**

Amplicon Processing

ASVs were processed using QIIME2 v2022.8 (1). Quality control on paired-end reads were conducted with FastQC v0.11.9 (2), followed by FIGARO (v1.0.0) to determine optimal trimming parameters for 16S reads. DADA2 v1.14.1 (3), implemented within QIIME2, was used to merge paired-end reads using FIGARO parameters and a maximum error rate of 0–2. Of note, the V4-V5 reads from the 18S rDNA region could not be merged, so V4 reads that didn't exceed a maximum error rate over 2 were used. Taxonomic classification was performed using the *Qiime2* plug-in, RESCRIPt (v2022.8.0). For 16S, a naive Bayes classifier trained on the SILVA database v138.1 (4) targeting the V4 region (~309 bp) was used. For 18S, classification targeted the V4–V5 region (~635 bp) using a Mothur-formatted PR2 database v5.1 (5, 6). For each cruise and rRNA type, three outputs were generated: ASV count tables, taxonomic assignments, and ASV sequences, which were imported into R for downstream analysis. In *R-studio* (v4.4.0, Puppy Cup), rarefaction curves were made to select appropriate sequencing depth thresholds (Table S6). Samples were rarefied with `rarefy_even_depth()` using *phyloseq* v1.44.0 (7).

Community Analysis

Microbial community analyses were conducted using the *vegan* v2.6-4 (8), *phyloseq* v1.44.0 (7), and *tidyverse* v2.0.0 (9) packages. We first calculated relative abundances for all ASVs by dividing ASV count observations by pre-determined rarefy thresholds (Table S6). Bray-Curtis-based Non-metric Multidimensional Scaling (NMDS) was performed using *vegan* v2.6-4. Post-hoc fitting of environmental factors (e.g. filter-size, region) and continuous variables (e.g. taxonomic groups) was performed on the NMDS ordination using the 'envfit' function to quantify how filter size, latitude, and phyla-level taxonomy explained compositional variance ($R^2$) of samples.

ASVs representing the collective community were categorized as either *persistent* (present in all three cruises) or *ephemeral* (present in only one or two cruises). To quantify how distinct each category was from the total community (persistent + ephemeral), Bray-Curtis dissimilarities were calculated between ephemeral vs. collective and persistent vs. collective groups. For each sample, ASV-level relative percent abundances were summed by phytoplankton group, and a pseudocount of 0.0001 was added to all values to avoid zeros. Dissimilarity was computed using vegdist(method = "bray").

Physio- and Biogeochemical Data Integrations

Physiochemical measurements (ie. seawater surface temperature and seawater surface salinity) and biogeochemical measurements (particulate organic carbon (POC), particulate organic nitrogen (PON), net community production (NCP) were obtained for the Simons Collaborative Marine Atlas Project (CMAP; https://simonscmap.com/; (10)) and correspond to data from Juranek *et al.* (11). Data were downloaded from the following links.

G1 NCP: https://simonscmap.com/catalog/datasets/KOK1606_Gradients1_Surface_O2Ar_NCP
G2 NCP: https://simonscmap.com/catalog/datasets/MGL1704_Gradients2_Surface_O2Ar_NCP
G3 NCP: https://simonscmap.com/catalog/datasets/KM1906_Gradients3_Surface_O2Ar_NCP

G1 POC/N: https://simonscmap.com/catalog/datasets/Gradients1_KOK1606_PPPCPN_UW

G2 POC/N: https://simonscmap.com/catalog/datasets/Gradients2_MGL1704_PPPCPN_UW
G3 POC/N: https://simonscmap.com/catalog/datasets/Gradients3_KM1906_PCPN_UW

Amplicon libraries were paired with corresponding POC, PON or NCP data using a collection time cutoff of 12 hours and location cutoff of 0.5 degrees of latitude. Only those amplicon libraries that could be linked to at least one type of data were used for subsequent analyses. Using this criterion, all 48 samples from 2016 had matched NCP data, while 38 had matched POC and PON data. In 2017, 56 of 72 samples had matched NCP, and 60 had matched POC and PON data. In 2019, 168 of 192 samples had matched NCP, while all had matched POC and PON data.

### Multi-level Mixed Modeling

To determine which phytoplankton groups best explained POC, PON, and NCP levels, we performed multivariate linear mixed modeling (MLMM) using the mmer(..., getPEV = TRUE, rcov = ~units) command within the *sommer* package v4.3.3 (12). ASV relative percent abundance was used to construct dissimilarity-based G-matrices for each phytoplankton group: persistent members housed within *Prochlorococcus*, *Synechococcus*, Archaeplastida, Haptophyta, Dinoflagellata within the Alveolata, and Stramenopiles belonging to Dictyochophyceae, diatoms (Coscinodiscophyceae, Bacillariophyceae, and Mediophyceae), Chrysophyceae, Pelagophyceae, Bolidophyceae, and Pinguiophyceae classes. To do this, we first added a pseudo-count of 1e-6 to all count observations. A center-log ratio (clr) was applied to each dataset, after which all data were merged to one master dataframe. Persistent phytoplankton taxa were retained while other ASVs were removed. For each phytoplankton group, a sample-by-ASV matrix was constructed from the master dataframe to calculate pairwise Euclidean distances which yielded a dissimilarity matrix per group. To avoid computational instability from zero distances, a small constant (0.001) was added to the diagonal - a regularization step commonly used to stabilize covariance matrix estimation in mixed modeling (13, 14).

We used the following equation to model the relationship between microbial community structure and biogeochemical variability across the North Pacific transect.

(a) $Y = \beta_0 + \beta_1 X_1 + \beta_2 X_2 + \beta_3 X_3 + Z_1\gamma_1 + Z_2\gamma_2 + Z_3\gamma_3 + Z_4\gamma_4 + Z_5\gamma_5 + Z_6\gamma_6 + \varepsilon$

(b) Feature ∼ time + filter + depth +
(1 | Archaeplastida) + (1 | Haptophyta) + (1 | Dinoflagellata) +
(1 | Stramenopiles) + (1 | Prochlorococcus) + (1 | Synechococcus)

where the Y represents the response variable (either POC, PON, or NCP), modeled individually and Box-Cox transformed using the bc_transform function from the bestNormalize package. $X_1$, $X_2$, $X_3$ correspond to fixed effects: time (Cruise × Month), filter (size fraction), and depth (binned into 0–15 m, 45–75 m, and 90–125 m ranges). $Z_n\gamma_n$ are the random effects representing dissimilarity-based G-matrices for each major phytoplankton group: Archaeplastida, Haptophyta, Dinoflagellata, Stramenopiles, *Prochlorococcus*, and *Synechococcus*. ε is the residual error term. Fixed effects were selected based on AIC and BIC minimization after testing multiple combinations. Model convergence was assessed using the summary() function in the sommer package, which reports convergence status and AIC/BIC values. For each model, we further examined residual distributions to confirm model assumptions. Residuals and fitted values were extracted and visualized using custom plotting functions that generated residuals vs. fitted value plots (to assess homoscedasticity), histograms with Shapiro-Wilk tests (to evaluate normality), and quantile-quantile (QQ) plots of residuals to assess distributional fit and model adequacy.

Network Construction and Analysis

Weighted gene co-expression network analysis (WGCNA v1.72-5 (15) was applied to persistent ASVs from phytoplankton-containing taxa and Box-Cox–transformed NCP , POC, and PON values to identify ASV modules correlated with these biochemical variables. A soft-thresholding power of 6 was selected using *pickSoftThreshold* (scale-free topology fit > 0.9; mean connectivity > 0). Networks were constructed with *blockwiseModules* (TOMType = "signed", deepSplit = 4, minModuleSize = 10, mergeCutHeight = 0.25) using a biweight midcorrelation (bicor) matrix to reduce outlier effects.

ASVs were hierarchically clustered based on relative percent abundance to define module memberships, and eigengenes were computed with *moduleEigengenes*. Module-trait correlations with NCP, POC, and PON were assessed using Pearson correlation, where correlations reflect statistical associations but not causality. Module composition was summarized by tallying ASVs per phytoplankton group, including species-level counts for Archaeplastida. Network structure was exported to *Cytoscape* v3.10.2 (16) for visualization and analysis using the ‘Network Analyzer’ tool. Additional soft-thresholding tests (powers 3–8) were run to evaluate the consistency of Archaeplastida presence in modules positively associated with NCP, POC, and PON across different network topologies.

Sparse Inverse Covariance Estimation for Ecological Association Inference (SpiecEasi) v1.1.0 (17) was used to infer direct associations between specific persistent phytoplankton and prokaryotic ASVs (Dataset S2). SpiecEasi networks were generated using the Meinshausen-Bühlmann (MB) method with lambda.min.ratio = 0.01 and nlambda = 20. The Stability Approach to Regularization Selection (StARS) with 50 subsampling iterations was used to select optimal sparsity parameters. Edges representing strong positive or negative associations (> +0.1 or < –0.1) were retained for visualization. To assess spatial relevance, we examined the latitude-specific distributions of Archaeplastida ASVs (identified in MLMM and WGCNA) and their first neighbors in the SpiecEasi network.

Supplemental References

1. Bolyen E, Rideout JR, Dillon MR, Bokulich NA, Abnet CC, Al-Ghalith GA, Alexander H, Alm EJ, Arumugam M, Asnicar F, Bai Y, Bisanz JE, Bittinger K, Brejnrod A, Brislawn CJ, Brown CT, Callahan BJ, Caraballo-Rodríguez AM, Chase J, Cope EK, Da Silva R, Diener C, Dorrestein PC, Douglas GM, Durall DM, Duvallet C, Edwardson CF, Ernst M, Estaki M, Fouquier J, Gauglitz JM, Gibbons SM, Gibson DL, Gonzalez A, Gorlick K, Guo J, Hillmann B, Holmes S, Holste H, Huttenhower C, Huttley GA, Janssen S, Jarmusch AK, Jiang L, Kaehler BD, Kang KB, Keefe CR, Keim P, Kelley ST, Knights D, Koester I, Kosciolek T, Kreps J, Langille MGI, Lee J, Ley R, Liu Y-X, Loftfield E, Lozupone C, Maher M, Marotz C, Martin BD, McDonald D, McIver LJ, Melnik AV, Metcalf JL, Morgan SC, Morton JT, Naimey AT, Navas-Molina JA, Nothias LF, Orchanian SB, Pearson T, Peoples SL, Petras D, Preuss ML, Pruesse E, Rasmussen LB, Rivers A, Robeson

MS, Rosenthal P, Segata N, Shaffer M, Shiffer A, Sinha R, Song SJ, Spear JR, Swafford AD, Thompson LR, Torres PJ, Trinh P, Tripathi A, Turnbaugh PJ, Ul-Hasan S, van der Hooft JJJ, Vargas F, Vázquez-Baeza Y, Vogtmann E, von Hippel M, Walters W, Wan Y, Wang M, Warren J, Weber KC, Williamson CHD, Willis AD, Xu ZZ, Zaneveld JR, Zhang Y, Zhu Q, Knight R, Caporaso JG. 2019. Author Correction: Reproducible, interactive, scalable and extensible microbiome data science using QIIME 2. Nat Biotechnol 37:1091–1091.

2. Andrews S. Babraham Bioinformatics - FastQC A Quality Control tool for High Throughput Sequence Data. https://www.bioinformatics.babraham.ac.uk/projects/fastqc/. Retrieved 28 March 2025.

3. Callahan BJ, McMurdie PJ, Rosen MJ, Han AW, Johnson AJA, Holmes SP. 2016. DADA2: High resolution sample inference from Illumina amplicon data. Nat Methods 13:581–583.

4. Quast C, Pruesse E, Yilmaz P, Gerken J, Schweer T, Yarza P, Peplies J, Glöckner FO. 2013. The SILVA ribosomal RNA gene database project: improved data processing and web-based tools. Nucleic Acids Res 41:D590–D596.

5. Schloss PD, Westcott SL, Ryabin T, Hall JR, Hartmann M, Hollister EB, Lesniewski RA, Oakley BB, Parks DH, Robinson CJ, Sahl JW, Stres B, Thallinger GG, Van Horn DJ, Weber CF. 2009. Introducing mothur: Open-Source, Platform-Independent, Community-Supported Software for Describing and Comparing Microbial Communities. Appl Environ Microbiol 75:7537–7541.

6. Guillou L, Bachar D, Audic S, Bass D, Berney C, Bittner L, Boutte C, Burgaud G, De Vargas C, Decelle J, Del Campo J, Dolan JR, Dunthorn M, Edvardsen B, Holzmann M, Kooistra WHCF, Lara E, Le Bescot N, Logares R, Mahé F, Massana R, Montresor M, Morard R, Not F, Pawlowski J, Probert I, Sauvadet A-L, Siano R, Stoeck T, Vaulot D, Zimmermann P, Christen R. 2012. The Protist Ribosomal Reference database (PR2): a catalog of unicellular eukaryote Small Sub-Unit rRNA sequences with curated taxonomy. Nucleic Acids Res 41:D597–D604.

7. McMurdie PJ, Holmes S. 2013. phyloseq: An R Package for Reproducible Interactive Analysis and Graphics of Microbiome Census Data. PLoS ONE 8:e61217.

8. Oksanen J, Simpson GL, Blanchet FG, Kindt R, Legendre P, Minchin PR, O'Hara RB, Solymos P, Stevens MHH, Szoecs E, Wagner H, Barbour M, Bedward M, Bolker B, Borcard D, Carvalho G, Chirico M, Caceres MD, Durand S, Evangelista HBA, FitzJohn R, Friendly M, Furneaux B, Hannigan G, Hill MO, Lahti L, McGlinn D, Ouellette M-H, Cunha ER, Smith T, Stier A, Braak CJFT, Weedon J, Borman T. 2025. vegan: Community Ecology Package (2.6-10).

9. Wickham H, Averick M, Bryan J, Chang W, McGowan LD, François R, Grolemund G, Hayes A, Henry L, Hester J, Kuhn M, Pedersen TL, Miller E, Bache SM, Müller K, Ooms J, Robinson D, Seidel DP, Spinu V, Takahashi K, Vaughan D, Wilke C, Woo K, Yutani H. 2019. Welcome to the Tidyverse. J Open Source Softw 4:1686.

10. Ashkezari MD, Hagen NR, Denholtz M, Neang A, Burns TC, Morales RL, Lee CP, Hill CN, Armbrust EV. 2021. Simons Collaborative Marine Atlas Project (Simons CMAP): An open-source portal to share, visualize, and analyze ocean data. Limnol Oceanogr Methods 19:488–496.

11. Juranek LW, White AE, Dugenne M, Henderikx Freitas F, Dutkiewicz S, Ribalet F, Ferrón S, Armbrust EV, Karl DM. 2020. The Importance of the Phytoplankton "Middle Class" to Ocean Net Community Production. Glob Biogeochem Cycles 34:e2020GB006702.

12. Covarrubias-Pazaran G. 2016. Genome-Assisted Prediction of Quantitative Traits Using the R Package sommer. PLOS ONE 11:e0156744.

13. Speed D, Balding DJ. 2014. MultiBLUP: improved SNP-based prediction for complex traits. Genome Res 24:1550–1557.

14. Bickel PJ, Levina E. 2008. Regularized estimation of large covariance matrices. Ann Stat 36:199–227.

15. Langfelder P, Horvath S. 2008. WGCNA: an R package for weighted correlation network analysis. BMC Bioinformatics 9:559.

16. Shannon P, Markiel A, Ozier O, Baliga NS, Wang JT, Ramage D, Amin N, Schwikowski B, Ideker T. 2003. Cytoscape: A Software Environment for Integrated Models of Biomolecular Interaction Networks. Genome Res 13:2498–2504.

17. Kurtz ZD, Müller CL, Miraldi ER, Littman DR, Blaser MJ, Bonneau RA. 2015. Sparse and Compositionally Robust Inference of Microbial Ecological Networks. PLOS Comput Biol 11:e1004226.
